# Subtilases turn on pectin methylesterase activity for a robust apoplastic immunity

**DOI:** 10.1101/2022.07.28.501549

**Authors:** Daniele Coculo, Daniele Del Corpo, Miguel Ozáez Martínez, Pablo Vera, Gabriella Piro, Monica De Caroli, Vincenzo Lionetti

## Abstract

Plants involve a fine modulation of pectin methylesterase (PME) activity against microbes. PME activity can promote the cell wall stiffening and the production of damage signals able to induce defense responses. However, to date, the knowledge about the molecular mechanisms triggering PME activity during disease remains largely unknown. In this study, we explored the role of subtilases (SBTs), serine proteases consisting of 56 isoforms in *Arabidopsis thaliana*, as activators of PME activity in plant immunity. By using biochemical and reverse genetic approaches, we found that SBT3.3 and SBT3.5 are required to control PME activity and resistance to the fungus *Botrytis cinerea. Arabidopsis sbt3.3 and sbt3.5* knockout mutants showed a reduced induction of PME activity and an increased susceptibility to *B. cinerea. SBT3.3* expression is controlled by the damage-associated molecular patterns Oligogalacturonides. The *SBT3.3* overexpression overactivates PME activity, but only during fungal infection, resulting in an increased expression of the defense-related genes and in an enhanced resistance to *B. cinerea*. We revealed that SBT3.3 and the Pro-PME17 isoforms are both secreted in the cell wall exploiting distinct protein secretion pathways and a different kinetic. Our findings point to SBTs as a mechanism to switch on PME activity and the related pectin integrity signaling to strengthen plant immunity against pests, in a timely manner to avoid the growth-defense trade-off.

**One sentence Summary:** Subtilases arm pectin methylesterase activity against pathogens to switch on pectin integrity signalling, reinforcing plant immunity and avoiding the growth-defense trade-offs

## Introduction

Plants continuously perceive attacks by microorganisms activating a timely and effective immune response. Part of this recognition is exerted by plasma membrane Pattern Recognition Receptors (PRRs), which can detect Microbe-, Pathogen-, or endogenous Damage Associated Molecular Patterns (MAMPs, PAMPs or DAMPs)(Zhou and Zhang, 2020). Pattern-triggered immunity (PTI) constitutes the first line of plant inducible immunity that restricts pathogen proliferation. Since the immunity efforts often result in a growth defense trade-offs, a precise regulation of immune pathways becomes critical to optimize fitness (Lionetti et al., 2010; Huot et al., 2014; Ha et al., 2021). A pivotal role in this homeostasis is played by proteolysis mediated by proteases, widely implicated in plant stresses, although their action mechanisms remain largely unknown (Schaller, 2004; Hou et al., 2018). In particular, Subtilases (SBTs, Pfam00082), serine peptidases belong to family S8, (MEROPS database; https://www.ebi.ac.uk/merops/cgi-bin/famsum?family=S8) plays key roles in defense responses against microbes, plant parasitism and adaptation to the environmental changes (Figueiredo et al., 2018; Schaller et al., 2018; Ogawa et al., 2021). In *A. thaliana*, 56 SBT isoforms are divided into 6 subfamilies: SBT1, SBT2, SBT3, SBT4, SBT5, SBT6 (Rautengarten et al., 2005).

Specific SBTs are targeted to the plant Cell Wall (CW), the foremost interface at which interactions between plants and microbes take place, to trigger specific protease-activated defense signaling (Doehlemann and Hemetsberger, 2013). In this compartment, SBTs can process larger precursor zymogens (Pro-Proteins), that require specific activation processes for enzyme maturation (Khan and James, 1998). SBT6.1 (S1P) can process the pro-peptide form of RAPID ALKALINIZATION FACTOR 23 (RALF23) modulating immune response against *Pseudomonas syringae* (Stegmann et al., 2017). In tomato, specific SBTs contributes to Jasmonic Acid-mediated resistance against *Manduca sexta* larvae, by processing the wound pro-hormone Prosystemin, releasing the bioactive peptide called Systemin (Meyer et al., 2016). The six genes of the tomato cluster SBT P69 (P69A–P69F) express proteins particularly involved in plant defense. P69B can cleave the *Phytophthora infestans* PC2, a cysteine-rich secreted apoplastic protein releasing immunogenic peptides that activates host immunity (Wang et al., 2021). In particular, P69C processes an extracellular matrix-associated Leucine-Rich Repeat protein (LRP), the first physiological target of an extracellular SBT identified in plants (Tornero et al., 1996). SBT3.3, the ortholog of tomato P69C in *Arabidopsis*, was discovered as regulator of *Arabidopsis* primed immunity (Ramirez et al., 2013). Although SBT3.3 was postulated to cleave an extracellular larger protein, the identification of its specific substrate and the mechanism by which SBT3.3 can trigger immunity remains to be identified.

During plant development, specific SBTs can process specific Pectin Methylesterases (PME; Pfam01095), enzymes regulating the methylesterification of Homogalacturonan (HG), the major constituent of pectin in the CW (Wormit and Usadel, 2018; Coculo and Lionetti, 2022). PMEs catalyse the hydrolysis of methylester bonds at C-6 of Galacturonic Acid (Gal A) residues, producing acidic pectins with negatively charged carboxyl groups, releasing methanol (MeOH) and protons into the apoplast. In *Arabidopsis*, 66 PME isoforms have been identified which can be classified into 2 groups on the basis of their structures (Pelloux et al., 2007; Dedeurwaerder et al., 2008). Group 1 comprises 21 PME isoforms, containing only the catalytic domain. The 45 PME isoforms of Group 2 (hereafter named Pro-PMEs) are organized as a polycistronic messenger RNA which resembles an operon-like gene cluster (Coculo and Lionetti, 2022). In Pro-PMEs, the PME catalytic domain is preceded by an N-terminal region (also referred as Pro region or PMEI-like domain) which shares structural similarities with functionally characterized PME Inhibitors (PMEIs) and that can acts as auto-inhibitor of PME activity (Bosch et al., 2005; Wolf et al., 2009; Del Corpo et al., 2020). This domain could impede a premature pectin de-methylesterification, during specific phases of their secretion, assembly, or modification contributing to a correct pectin functionality.

A local and strong induction of PME activity is exerted in *Arabidopsis thaliana* and other plants, when challenged with different microbial pathogens (Bethke et al., 2014; Lionetti, 2015; Lionetti et al., 2017). *Pro-PME17* isoform is strongly upregulated in response to several pathogens, defense hormones, elicitors or abiotic stress and its activity strongly contribute to the induction of defense-related PME activity (Bethke et al., 2014; Del Corpo et al., 2020). A finely tuned PME activity can play multiple functions in plant immunity. PMEs can produces de-methylesterified negatively charged carboxyl groups that can form consecutive calcium bonds with other HG molecules and thus strengthen the CW through the formation of “egg-boxes” structures (Senechal et al., 2014b; Del Corpo et al., 2020). Plant PME activity can also favour DAMP related immunity. In particular, the PME activity could favour the release and perception of de-methylesterified oligogalacturonides (OGs), the well characterized DAMP, locally released in cooperation with endo-polygalacturonases (PGs), and able to trigger plant immunity (Osorio et al., 2008; Osorio et al., 2011; Ferrari et al., 2013; Kohorn et al., 2014). OGs are perceived in *Arabidopsis* by the pathogen recognition receptor WALL-ASSOCIATED KINASE1 (WAK1) to activate plant immune responses (Brutus et al., 2010; Kohorn and Kohorn, 2012). WAK2 requires PME-mediated pectin de-methylesterification to activate OGs-dependent stress responses (Kohorn et al., 2014). Interestingly, WAK1, WAK2 and another PRR, FERONIA (FER), preferentially bind to de-methylesterified pectins (Decreux and Messiaen, 2005; Kohorn et al., 2014; Feng et al., 2018; Guo et al., 2018; Lin et al., 2021). The de-methylesterification of pectin by PMEs represents the main mechanism to generate plant-derived methanol (MeOH), a DAMP like alarm signal (Hann et al., 2014). OGs and MeOH are able to trigger defensive priming effect setting the stage for the intra and inter-plant immunity (Dorokhov et al., 2012; Komarova et al., 2014; Gamir et al., 2021; Giovannoni et al., 2021).

Precise switch on/off mechanisms regulating plant immunity are required because they must be timely triggered for an efficient immune response and they cannot be persistent given the well-known defense growth trade-off (He et al., 2022). The need of a post-transcriptional control of PME activity by specific PME inhibitors (PMEIs) late during plant invasion by microbes was previously demonstrated (An et al., 2008; Lionetti et al., 2017; Liu et al., 2018). The involvement of SBTs as posttranscriptional processing of Pro-PMEs and regulation of PME activity in plant immunity was still not investigated. Here, we found that all the PME isoforms induced in *A. thaliana* against the necrotrophic fungus *Botrytis cinerea* possess a Pro-region. Exploiting reverse genetic approaches and biochemistry, we demonstrated that SBT3.3 and SBT3.5 contribute to the induction of PME activity in *Arabidopsis* immunity against *Botrytis*. Notably, we found that SBT3.3 processes defense related Pro-PMEs as substrate and trigger specific defense related genes. SBT3.3 and Pro-PME17 follows distinct secretion pathways to reach the apoplast where they later colocalize. Our results provide strong evidence that the processing of Pro-PMEs by SBTs represents a mechanism to switch on PME activity and the related pectin integrity signaling for a timely immune response, avoiding the growth-defense trade-offs.

## RESULTS

### The processing of Pro-PME zymogens is required in *Arabidopsis-Botrytis* interaction

*PMEPCRA, PME3, PME17, PME20, PME21*, and *PME41* are the PME isoforms exhibiting a significant altered timing and level of expression in *A. thaliana* - *B. cinerea* interaction and in several other pathosystems (Lionetti et al., 2012; Bethke et al., 2014; Lionetti et al., 2017). Interestingly, after a careful comparison of the sequences we realized that all these isoforms are zymogens, expressed as Pro-PME inactive precursors (Supplemental Figure S1). This strongly suggests that a proteolytic cleavage, exerted by specific proteases, is requested for PME activation in *Arabidopsis* immunity against *Botrytis*. While the PME catalytic domain is highly conserved, we intriguingly noticed here a variability in the processing motif between the different PME isoforms. Pro-PMEPCRA, Pro-PME17 and Pro-PME21 showed a RKLL processing motif, while the RRKL motif characterized the Pro-PME20 and Pro-PME41 isoforms. Instead, two processing motifs, RKLK and RRLL were present in the Pro-PME3. Other motifs with the consensus sequence FPSW and LPLKMTERARAV are partially conserved between the isoforms. All these observations could imply the involvement of multiple proteases-PME pairs.

### *SBT3.3* and *SBT3.5* are induced in *Arabidopsis* during *B. cinerea* as part of PTI

We explored the possibility that SBTs are involved in the processing and activation of Pro-PMEs against microbes. Exploiting publicly available microarray data, we selected two genes, *SBT3.3* and *SBT3.5* showing a significative induction during *B. cinerea* infection (Supplemental Figure S2A). Interestingly, both genes are co-expressed with *Pro-PME17*, a functional PME involved in the resistance to *Botrytis cinerea* (Supplemental Figure S2B) (Lionetti et al., 2012; Del Corpo et al., 2020). To confirm the transcriptomic analysis and to gain more insight in the kinetic of expression of these genes during the disease, the expression levels of *SBT3.3* and *SBT3.5* were quantified at different times in *B. cinerea-infected* and Mock-inoculated *A. thaliana* wild type (WT) leaves by qPCR (Figure 1A and B). *SBT3.3* is expressed only in infected tissues while *SBT3.5* shows a basal expression also in Mock-inoculated leaves. *SBT3.3* and *SBT3.5* were significantly induced at 24 hpi and further increased at 48 hpi, when they showed about a significant 15- and 5-fold induction, respectively, in infected leaves compared to Mock inoculated tissues. These data clearly reveal an involvement of *SBT3.3* and *SBT3.5* in *Arabidopsis-Botrytis* interaction. Interestingly, the expression of both *SBTs* is also triggered in *Arabidopsis* against several pathogens and by different bacterial and fungal elicitors of plant defense responses (Supplemental Figure S3). We next investigated whether pectin-related DAMPs can impact on the expression of the selected SBTs. *Arabidopsis* WT seedlings were treated with OGs or water, and the expression of *SBT3.3* and *SBT3.5* was evaluated at 1 h post treatment. A 7-fold induction of *SBT3.3* was revealed after treatment with OGs (Supplemental Figure S3 and Figure 1C), while the expression of *SBT3.5* was not induced (Supplemental Figure S3 and Figure 1D). All these results indicate that *SBT3.3* and *SBT3.5* are part of *Arabidopsis* immune responses against *B. cinerea* and several pathogens.

**Figure 1.**
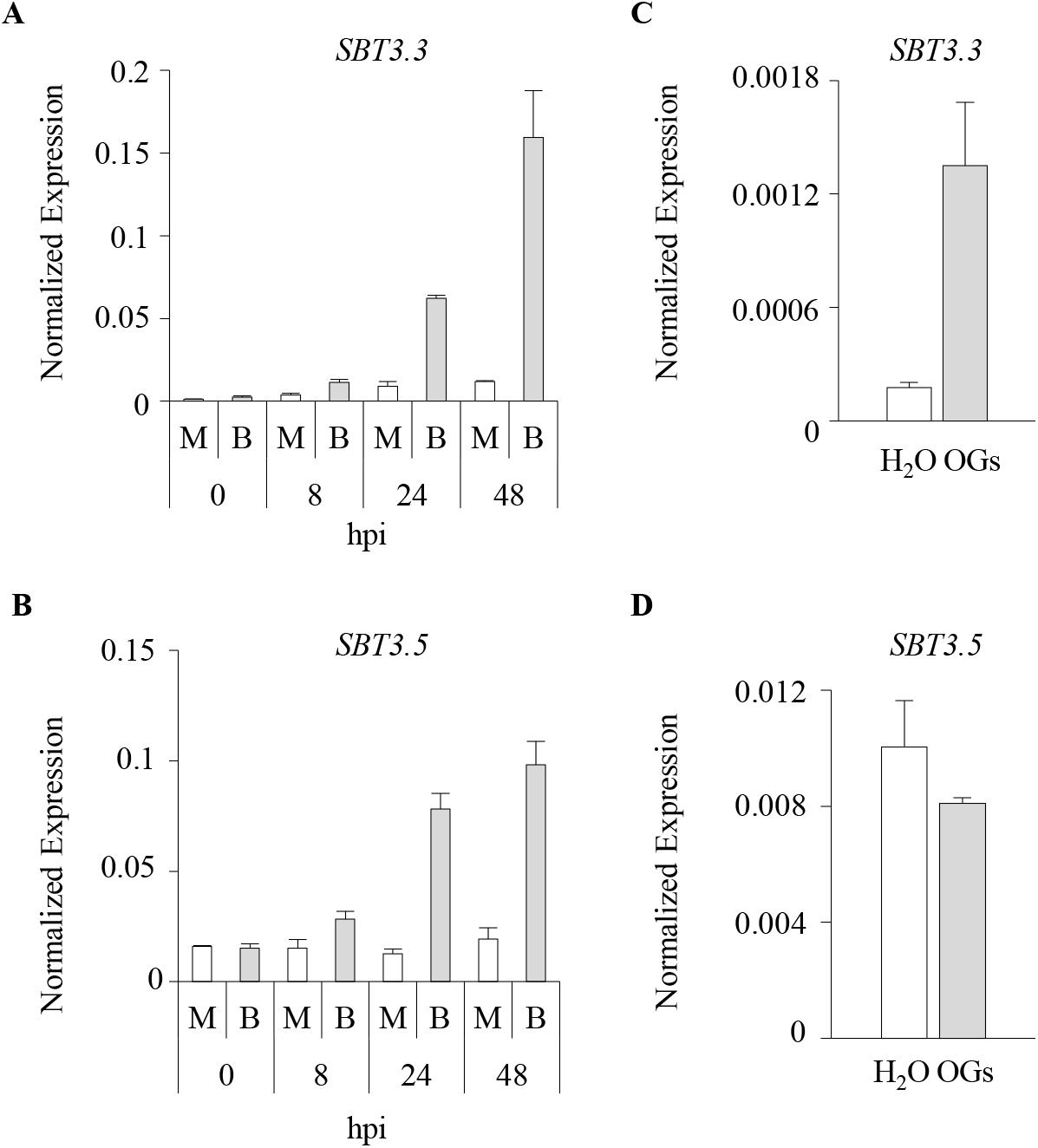
*SBT3.3* and *SBT3.5* are induced in *Arabidopsis* during *B. cinerea* infection and SBT3.3 is induced by OGs treatment. **A**) *SBT3.3* and **B)** *SBT3.5* expression was analysed in *B. cinerea* infected or mock-inoculated leaves at the indicated hours post inoculation (hpi) by qPCR. M = Mock inoculated leaves. B = *B. cinerea* inoculated leaves. The expression of **C)** *SBT3.3* and **D)** *SBT3.5* was analysed in Arabidopsis WT seedlings after 1 hour of treatment with OGs or with water (H_2_O) as a control. The expression levels were normalized to *Ubiquitin5* expression. Data represent mean ± SD (n=3). The experiments were repeated three times with similar results.

### SBT3.3 and SBT3.5 contribute to *Arabidopsis* PME activation and resistance to *B. cinerea*

To explore on the possibility that SBT3.3 and SBT3.5 may be involved in the activation of Pro-PMEs in plant immunity we exploited a reverse genetic approach. Two T-DNA insertional mutants for the selected genes were isolated (Figure 2A). The insertions were localized in the fifth and ninth exons, respectively, for *sbt3.3-1* (SALK_086092) and *sbt3.3-2* (SALK_107470.45.45) mutants and in the first exon and in the second intron, respectively for *sbt3.5-1* (SAIL_400_F09) and *sbt3.5-2* (GABI_672C08) mutants. The level of *SBT3.3* and *SBT3.5* expression was compared in leaves of mutants and WT plants after *B. cinerea* and Mock inoculation by qPCR (Figure 2B). All mutants fail to induce gene expression in fungus-challenged leaves, resulting in *sbt3.3* and *sbt3.5* KO lines. No significant defects were observed in the rosette growth and seed germination in *sbts* mutants which showed growth parameters similar to WT plants (Figure 2C and S4A and B). To understand if the lack of *SBTs* expression can influence pectin composition we compared the monosaccharide composition of matrix polysaccharides extracted from rosette leaves of *sbts* mutants and WT plants. All mutants analysed showed a molar percentage of monosaccharides not significantly different from that observed in WT plants (Figure 2D). After establishing that the two SBTs are involved in *Arabidopsis-Botrytis* interaction we evaluated their possible contribution in the control of PME activity during fungal infection. The level of PME activity was quantified in leaves of WT, *sbt3.3-1, sbt3.3-2, sbt3.5-1* and *sbt3.5-2* plants at 24 hours post *Botrytis* and Mock inoculation using histochemical and biochemical assays based on Ruthenium Red (RR), a cationic dye with six positive charges able to bonds with the acidic groups of HG (Lionetti, 2015)(Figure 3A). A significant lower induction of PME activity was observed in infected *sbt3.3-1, sbt3.3-2, sbt3.5-1* and *sbt3.5-2* mutants (respectively of 3.7-, 3.1, 1.9- and 1.8-fold reduction) compared to WT (Figure 3B). These results unveil for the first time that Pro-PMEs are substrates for SBT3.3 and indicate that SBT3.3 and SBT3.5 contribute to control the PME activity in *Arabidopsis* immunity to *Botrytis*. Interestingly, the significant difference found in *sbt3.3* respect to *sbt3.5* mutants suggest that the two proteases can process different Pro-PMEs isoforms during disease. No significant differences in PME activity were observed between Mock-inoculated WT plants and *sbt* mutants. This was expected for *sbt3.3* mutants, as *SBT3.3* is not basally expressed in leaves (Figure 1A). In *sbt3.5* mutants, the lack of the basal *SBT3.5* expression in Mock-inoculated leaves (Figure 2B) does not significantly affect the monosaccharide composition and the total PME activity in leaves (Figure 2D and 3B) as well as on the rosette growth and seed germination (Figure 2C and Supplemental Figure S4). Therefore, we evaluated the susceptibility of *sbt3.3* and *sbt3.5* mutants to *B. cinerea* (Figure 4A and B). The local symptoms of the fungus were significantly higher in both *sbt3.3* and *sbt3.5* mutants respect to WT. In particular, the lesion areas produced by the fungus were larger in *sbt3.3-1, sbt3.3-2, sbt3.5-1, sbt3.5-2* mutants (respectively, of 2.2-, 2.1, 1.7- and 1.7-folds) respect to WT. A greater development of *Botrytis* mycelium was detected in *sbt* mutants respect to WT plants using trypan blue staining (Figure 4B). All our results indicate that, although at different extent, both SBTs contribute to trigger PME activity and the resistance to *Botrytis*. A correlation between the level of resistance and the amount of PME activity expressed in the tissues were observed between the different genotypes analysed (Figure 3 and 4).

**Figure 2.**
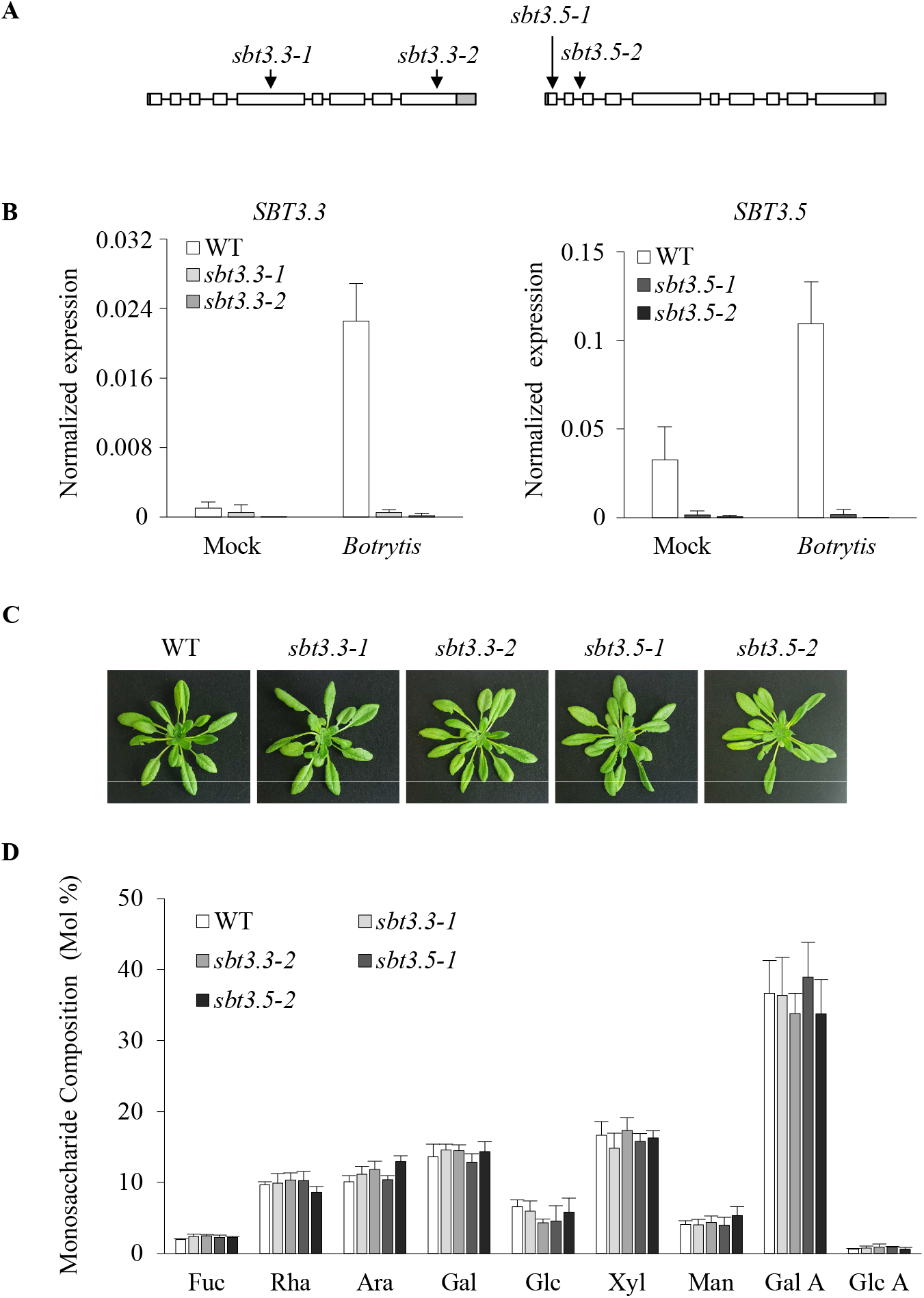
Isolation of Arabidopsis *SBT3.3* and *SBT3.5* T-DNA insertional mutants. **A)** Schematic representation of *SBT3.3* and *SBT3.5* gene structures. The localization of T-DNA insertions in the gene sequences are shown (black arrows). 5’-UTR and 3’-UTR are represented in grey, exons in white and introns as a black line. **B)** The expression of *SBT3.3* and *SBT3.5* was analyzed in the respective mutants and WT plants at 24-hour post inoculation with *B. cinerea*. The expression levels were normalized to *Ubiquitin 5* expression. The experiments were repeated three times with similar results. **C)** Representative pictures illustrating the morphology of vegetative rosettes of *sbt3.3* and *sbt3.5* mutants compared with WT. **D)** The monosaccharide composition of cell wall matrix polysaccharides was compared in leaves of *sbt3.3* and *sbt3.5* mutants and WT plants. The molar percentages of Fucose (Fuc), Rhamnose (Rha), Arabinose (Ara), Galactose (Gal), Glucose (Glu), Xylose (Xyl), Mannose (Man), Galacturonic Acid (Gal A) and Glucuronic Acid (Glc A) were quantified. The results represent the mean ± SD (n>3). Monosaccharide datasets are not significantly different according to analysis of variance (ANOVA) followed by Tukey’s test (P < 0.05).

**Figure 3.**
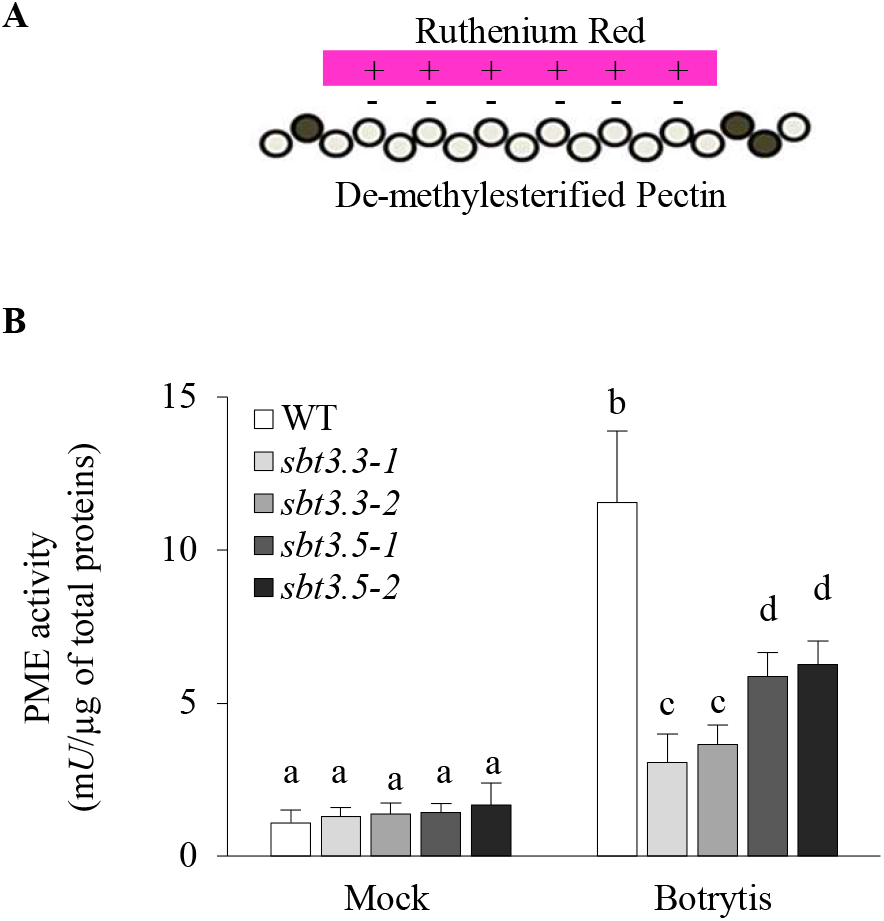
SBT3.3 and SBT3.5 regulate PME activity during disease. **A)** Schematic representation of the binding of Ruthenium Red to de-methylesterified pectin. **B)** Leaves of Arabidopsis WT and of *sbt* mutants were inoculated with *Botrytis* and Mock and PME activity were quantified at 24-hour post inoculation (hpi). The results represent the mean ± SD (n = 6). The different letters on the bars indicate datasets significantly different according to ANOVA followed by Tukey’s test (P < 0.05).

**Figure 4.**
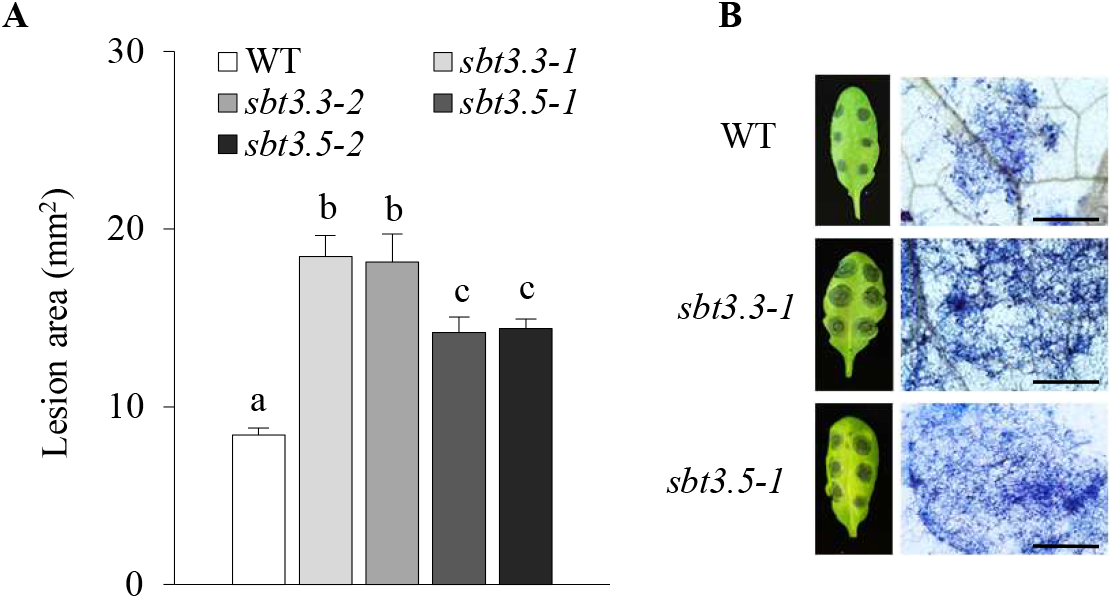
The lack of *SBT3.3 and SBT3.5* expression reduces *Arabidopsis* resistance to *Botrytis cinerea*. **A)** Quantification of lesion areas produced by the spreading of the fungus at 24 hours post infections. Data represent mean ± SE (n >14 lesions for each genotype). Different letters indicate statistically significant differences, according to ANOVA followed by Tukey’s significance test (P < 0.05). **B)** Photographs showing lesion areas produced by *Botrytis* on leaves of Arabidopsis WT, *sbt3.3- 1 and sbt3.5-1* mutants (left) and microphotographs showing *Botrytis* colonization revealed by trypan blue staining (right). Bars = 1mm.

### SBT3.3 overexpression overactivates defense related PME activity restricting *Botrytis* infection

Given the lower induction of PME activity observed in *sbt3.3* mutants compared to those of SBT3.5, we deepen our study on SBT3.3 exploring the effect of its overexpression in *Arabidopsis* on PME activity and *Botrytis* resistance. Two transgenic *Arabidopsis* lines, named *SBT3.3-OE1* and *SBT3.3- OE2* exhibiting high level of SBT3.3 expression in leaves were characterized (Figure 5A). Interestingly, an 85- and 90-fold overexpression was revealed, respectively, in leaves of *SBT3.3-OE1* and *SBT3.3-OE2* lines respect to WT leaves when infected with *B. cinerea*. No growth abnormalities were observed in *SBT3.3* overexpressing plants (Figure 5B and S4). The CW monosaccharide composition in rosette leaves of SBT3.3-OE lines is not significantly different from that quantified in the WT plants (Figure 5C). These results are consistent with those obtained with the mutants and corroborate the conclusion that the expression of *SBT3.3* is dedicated to the *Arabidopsis* immune response. The level of PME activity was quantified in leaves of WT, *SBT3.3OE-1 and SBT3.3OE-2* plants at 24h post *Botrytis-* and Mock-inoculation (Figure 6A). A significant higher induction of defense-related PME activity was observed in the infected *SBT3.3* overexpressing lines compared to control. *SBT3.3OE-1* and *SBT3.3OE-2* lines showed, respectively, a 3.1- and 2.3- fold increase of PME activity in the *B. cinerea* infected leaves compared to WT. Despite the *SBT3.3* overexpression in transgenic Mock-inoculated plants, the level of PME activity were not significantly different from that measured in WT plants (Figure 6A). This result highlights that SBT3.3 triggers PME activity exclusively in *Arabidopsis* defense against the fungus. The levels of methylesters remained in the CW after 24 h of *B. cinerea* inoculation were compared between *SBT3.3-OE* lines and WT plants. A significant lower amount of methylesters was revealed in the CW extracted from *B. cinerea* infected leaves of *SBT3-3-OE1* and *SBT3.3-OE2* lines respect the WT plants (Figure 6B). This result confirms that SBT3.3 processes the Pro-PMEs in plant defense and that a higher PME activity was released by SBT3.3 overexpression in transgenic plants challenged with the fungus respect the control.

**Figure 5.**
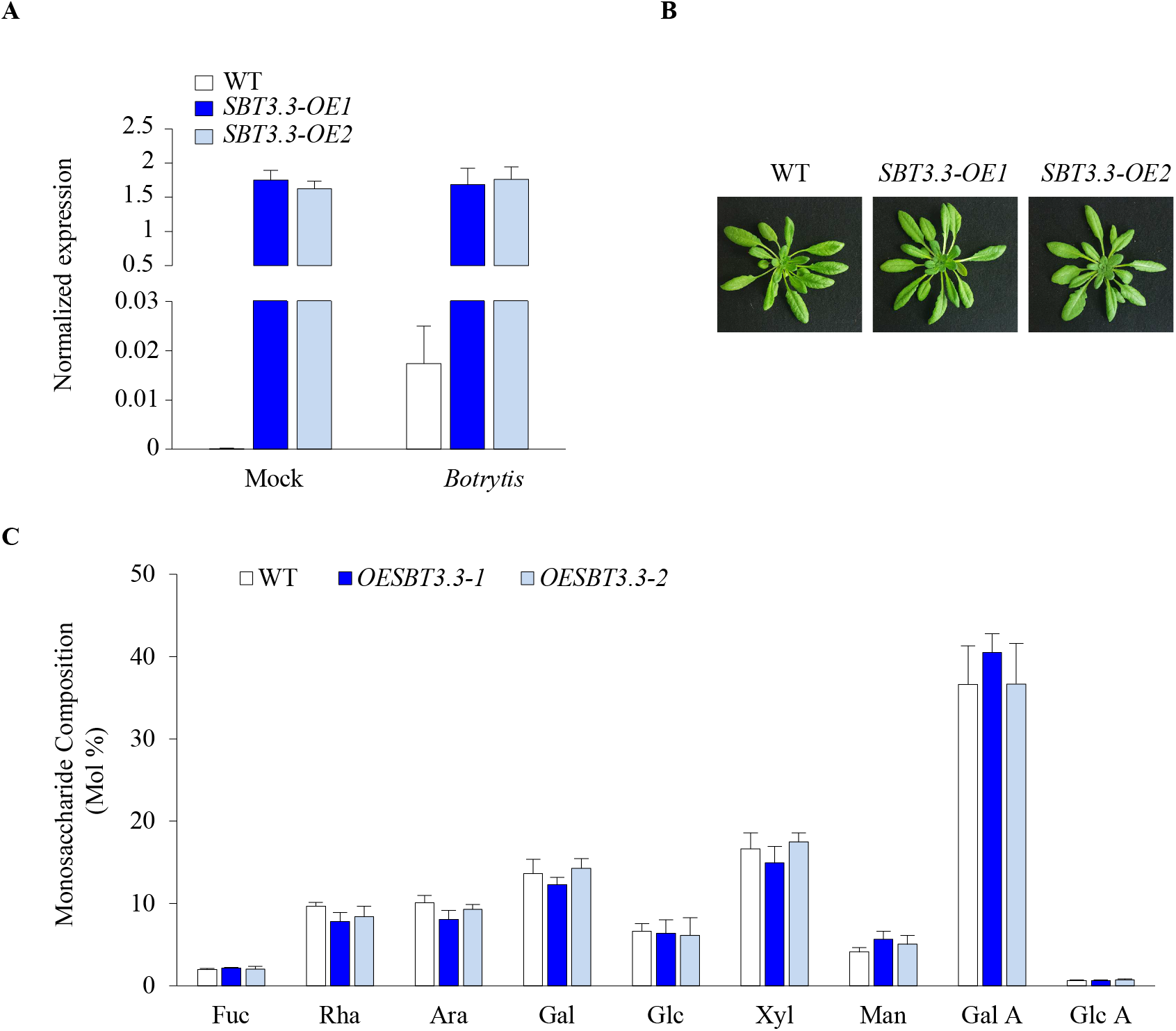
The *SBT3.3* overexpression in Arabidopsis does not alter rosette growth and cell wall composition. **A)** The expression of *SBT3.3* was analyzed in the *SBT3.3-OE1* and *SBT3.3-OE2* transgenic plants at 24-hour post inoculation with Mock and *B. cinerea* spores. The expression levels were normalized to *Ubiquitin5* expression. The experiments were repeated three times with similar results. **B)** Representative pictures illustrating the morphology of vegetative rosettes of *SBT3.3* overexpressing plants compared with WT plants. **C)** The molar percentages of Fucose (Fuc), Rhamnose (Rha), Arabinose (Ara), Galactose (Gal), Glucose (Glu), Xylose (Xyl), Mannose (Man), Galacturonic Acid (Gal A) and Glucuronic Acid (Glc A) were quantified. The results represent the mean ± SD (n > 3). Monosaccharide datasets are not significantly different according to analysis of variance (ANOVA) followed by Tukey’s test (P < 0.05).

**Figure 6.**
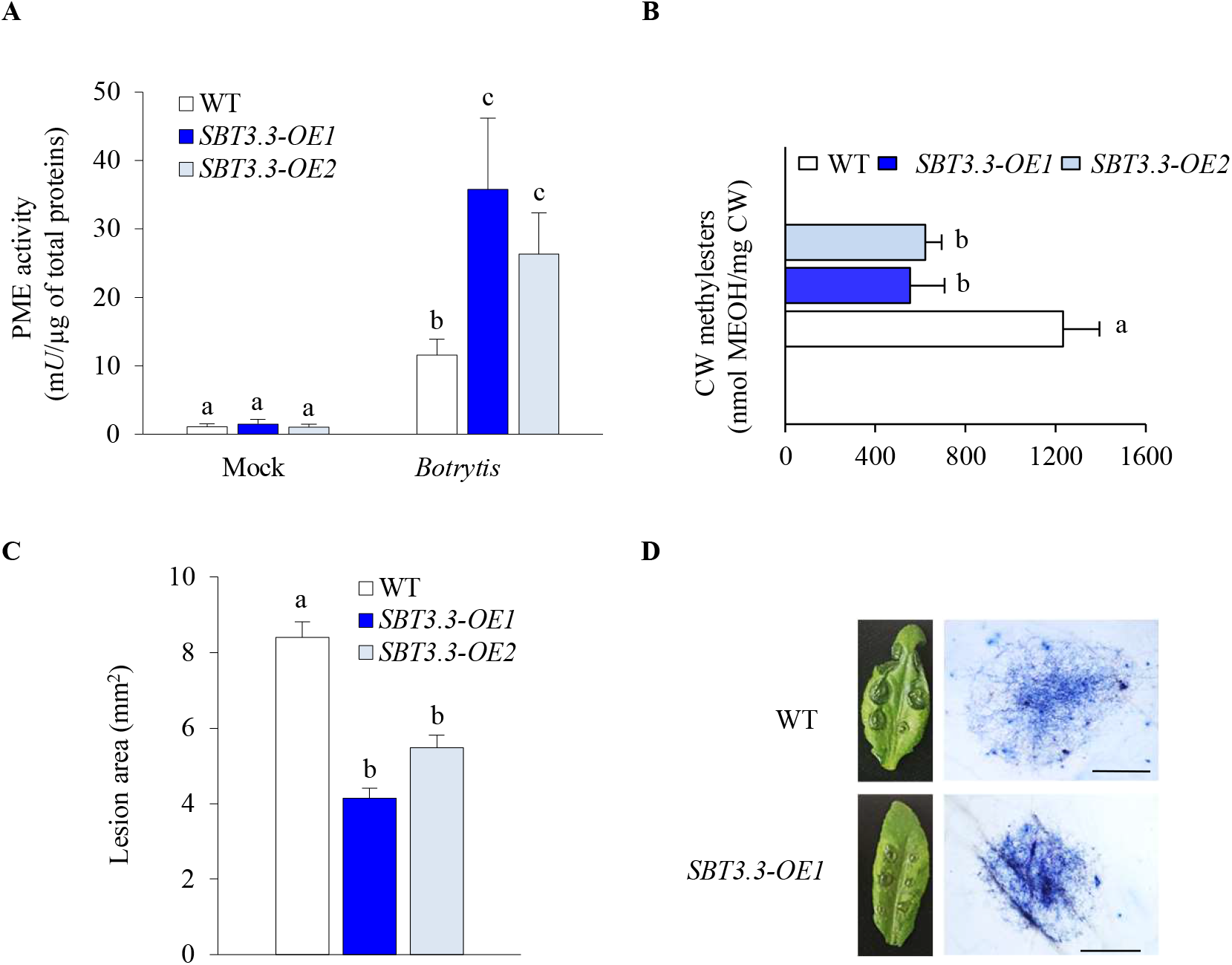
The *SBT3.3* overexpression overactivates PME activity restricting *Botrytis* infection. **A)** Leaves of Arabidopsis WT and of *SBT3.3-OE* plants were inoculated with *Botrytis* and Mock and PME activity were quantified at 24-hour post inoculation. The results represent the mean ± SD (n = 6). **B)** The amount of methylesters in the cell wall were quantified at 24-hour post inoculation. The results represent the mean ± SD (n = 6). **C)** Quantification of lesion areas produced by the spreading of the fungus at 24 hours post infections. Data represent mean ± SE (n>15 lesions for each genotype). **D)** Photographs showing lesion areas produced by *Botrytis* on leaves of WT and *SBT3.3-OE1* plants (left) and microphotographs showing *Botrytis* colonization revealed by trypan blue staining. Bars = 1mm. The different letters in A, B and C indicate datasets significantly different according to ANOVA followed by Tukey’s test (P < 0.05).

Next, we evaluated the susceptibility of *SBT3.3-OE* lines to *B. cinerea*. The local symptoms of the fungus were significantly lower in both *SBT3.3-OE* plants when compared with the WT (Figure 6C and D). In particular, the lesion area produced by the fungus in *SBT3.3-OE1* and *SBT3.3-OE2* mutants are 2- and 1.5-fold lower compared to WT. A minor development of *Botrytis* mycelium was detected in *SBT3.3-OE1* plants respect to WT plants using trypan blue staining (Figure 6D). All these results support an important role of SBT3.3 as trigger of PME activity in *Arabidopsis* immunity to *Botrytis* and indicate that the SBT3.3 overexpression confers to *Arabidopsis* a higher resistance to *B. cinerea* respect to WT.

### *SBT3.3* overexpression enhances immune responses against *Botrytis*

The next goal was to verify whether the improved resistance observed in the SBT3.3 overexpressing lines could be linked to a more robust immune response. The induction of an array of defense related genes important for *Arabidopsis* defense against *Botrytis* was compared between WT and *SBT3.3-OE1* plants inoculated with Mock and *Botrytis. WRKY33* is a key transcriptional regulator of hormonal and metabolic responses toward *B. cinerea* infection (Birkenbihl et al., 2012). *WRKY33* can targets the promoter of *PHYTOALEXIN DEFICIENT3 (PAD3)* encoding an enzyme required for the synthesis of antimicrobial camalexin (Zhou et al., 1999). The cytochrome P450 enzyme *CYP81F2* is essential for the pathogen-induced accumulation of 4-methoxy-indol-3-ylmethyl Glucosinolate (Clay et al., 2009; AbuQamar et al., 2017). *WAK2* is a wall associated kinase mediating pectin related immunity and activated by de-methyl esterified pectin to trigger defense responses (Kohorn et al., 2014). *PME17* and *PMEI10* are two regulators of PME activity in plant immunity to *B.cinerea* (Lionetti et al., 2017; Del Corpo et al., 2020). Interestingly, a higher induction of *WRKY33, PAD3*, and *WAK2* were observed in *SBT3.3-OE1* plants respect the WT only when challenged with the fungus (Figure 7). *CYP81F2* showed levels of expression similar between WT and *SBT3.3-OE1* plants. However, considering the lower level of *B. cinerea β-tubulin* and the lower fungal symptoms observed in transgenic plants (Figure 7 and Figure 6C and D) it is possible to conclude that also CYP81F2 expression is stimulated by *SBT3.3* overxpression. The *Pro-PME17* and *PMEI10* expressions seems independent from SBT3.3 activity since their inductions follow the extent of fungal infection and *B. cinerea β-tubulin* expression. These findings indicate that SBT3.3 contribute to enhance *Arabidopsis* immunity, most likely, improving the production of antimicrobial compounds and reinforcing cell wall integrity surveillance.

**Figure 7.**
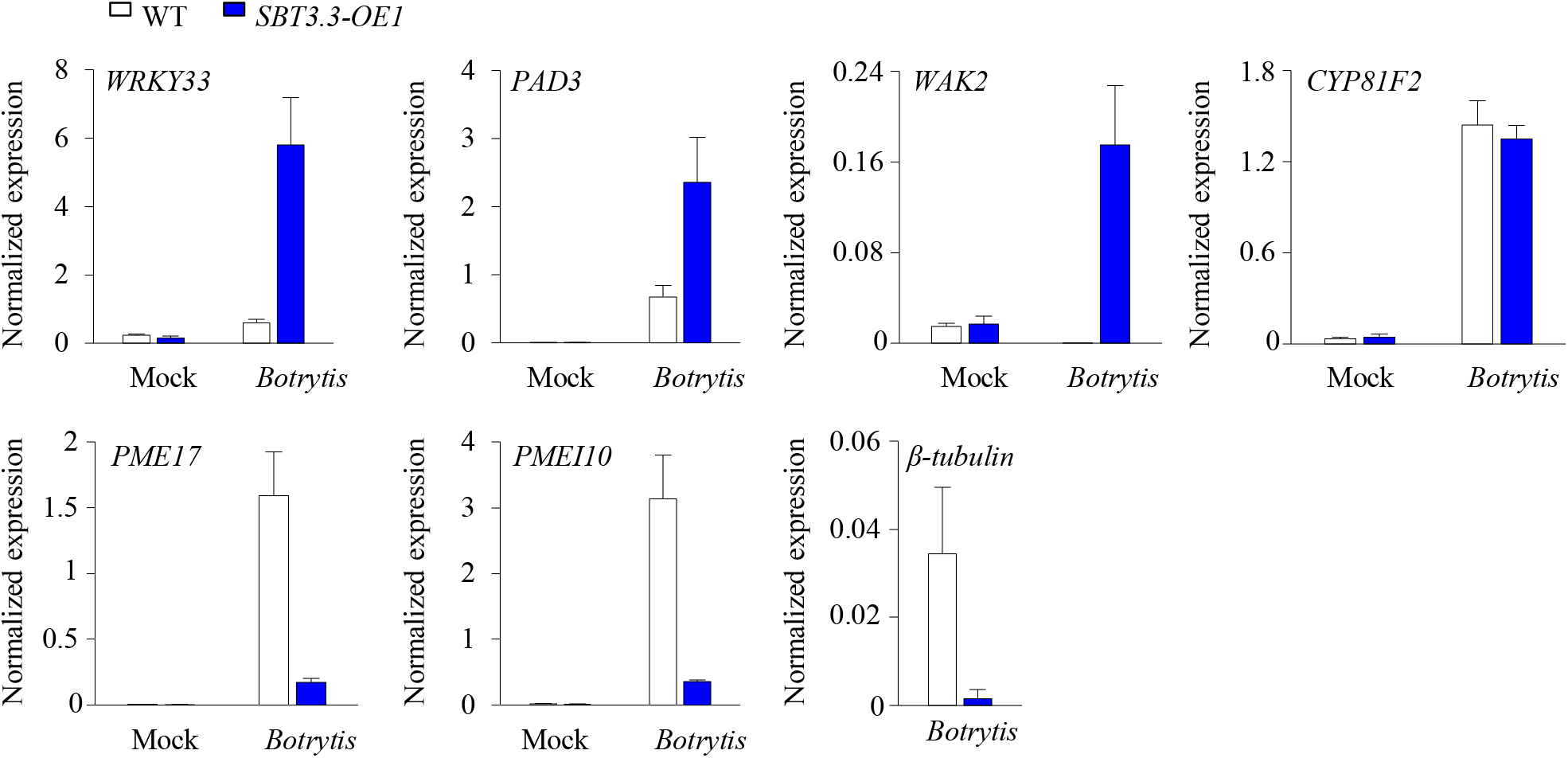
*SBT3.3* overexpression enhances immune responces. The induction of the expression of defence genes *WRKY33, PAD3, WAK2, CYP81F2, PME17, PMEI10* were compared between *SBT3.3- OE1* line and WT plants challenged with *Botrytis*. The mRNA expression was analysed at 24 hpi in *B. cinerea-* or mock-inoculated leaves by qPCR. *Botrytis β-tubulin* expression was also quantified. The expression levels were normalized to *Ubiquitin5* expression. The experiments were repeated three times with similar results.

We next explored if SBTs expression can influence Hydrogen Peroxide production, an important defense response to *Botrytis* infestation (Pogorelko et al., 2013; Siegmund and Viefhues, 2016). Leaves of WT, *sbt3.3-1, sbt3.5-1* and *SBT3.3-OE1* plants, were inoculated with Mock or *B. cinerea* and stained with 3,3’-diaminobenzidine, a chromogen able to reveal the H2O2 accumulation in the tissue (Reem et al., 2016). No accumulation of H2O2 was observed in Mock-inoculated leaves of all genotypes analyzed and a similar DAB staining was revealed in all the leaves challenged with *Botrytis*. These results indicate that H2O2 production is not dependent from SBTs mediated signaling (Supplemental Figure S5).

### SBT3.3 and Pro-PME17 are secreted through distinct protein secretion pathways in the apoplast where lastly colocalized

SBT3.3 was previously found in the extracellular compartment, but its secretion pathway was never explored (Ramirez et al., 2013). Moreover, Pro-PME17 was predicted to be an apoplastic protein, but its localization was never demonstrated. With the aim to deepen our knowledge on the secretion patterns and the final subcellular destination of the two proteins, a fluorescent variant of SBT3.3 and of Pro-PME17 were constructed by fusing the respective encoding cDNA to *yfp* and *gfp*, under the control of *35S* promoters. The fluorescent tag was linked to the C-terminus of both proteins, as none of them are predicted to have the ω site of the glycosylphosphatidylinositol (GPI) anchor addition (predGPI software; http://gpcr2.biocomp.unibo.it/predgpi/), as previously found for AtPMEI1 (De Caroli et al., 2011). *SBT3.3-YFP* and *Pro-PME17-GFP* constructs were transiently expressed in the epidermal cells of tobacco leaves and observed by laser scanning confocal microscopy (Figure 8). Both proteins accumulate in the apoplast as confirmed by colocalization with the CW marker XTH11-RFP (De Caroli et al., 2021) (Figure 8A-C, Q-S) and plasmolysis of epidermal cells co-expressing both *pm-rk*, marker of plasma membrane (Nelson et al., 2007) and *SBT3.3-YFP* or *Pro-PME17-GFP* (Figure 8D, T). Nevertheless, the fluorescence signals relative to SBT3.3-YFP and Pro-PME17-GFP showed different secretion patterns. SBT3.3-YFP exhibited a typical cytosolic fluorescent pattern, labelling the nucleoplasm and appearing excluded by the ER and Golgi stacks, without any colocalization with the specific organelle markers, RFP-HDEL and ST52-mCherry, respectively (Nelson et al., 2007) (Figure 8A-J). Other than nucleoplasm and cell wall, SBT3.3-YFP labelled small cytosolic punctate structures that significantly colocalized with the Exocyst Positive Organelles (EXPOs) marker Exo70E2-GFP (Wang et al., 2010; Wang et al., 2017) indicating the possible involvement of EXPOs in an unconventional trafficking of the chimera toward the CW (Figure 8K-M). Proteins that follow the Unconventional Secretion Pathway (UPS) bypass the Golgi and are insensitive to brefeldin A (BFA), an inhibitor of ER-Golgi trafficking (Zhang et al., 2011). BFA treatment did not affect SBT3.3-YFP secretion pattern while clearly perturbed the Golgi organization labelled by ST52-mCherry (Figure 8N-P). A conventional secretion pathway is followed by Pro-PME17-GFP that labelled the ER, continuous to the nuclear envelope, and small punctate structures colocalizing with the ER and the Golgi markers, respectively (Figure 8U-Z). Intriguingly, the Pro-PME17 reaches the apoplast before the SBT3.3. At 48 h post transformation, the CW was intensely labelled by Pro-PME17-GFP and, at approximately 60 h by SBT3.3-YFP protein fusion (Figure 8A and Q). These results indicate that both SBT3.3 and Pro-PME17 are correctly secreted into the apoplast by using different secretion pathways and in a temporally staggered manner. We further explored the timing of SBT3.3 and Pro-PME17 colocalization into the cell wall using a new version of *Pro-PME17* tagged with *rfp*, under the control of *35S* promoter transiently co-expressed with *SBT3.3-YFP* in the epidermal cells of tobacco leaves. We confirmed that the the Pro-PME17 and SBT3.3 colocalize at the CW, and this is favoured over time as revealed by colocalization parameters (Figure 9). All these results strongly support the hypothesis that Pro-PME17 is secreted in an unprocessed and inactive form in the CW where, later when required, is activated by SBT3.3 to trigger immunity.

**Figure 8.**
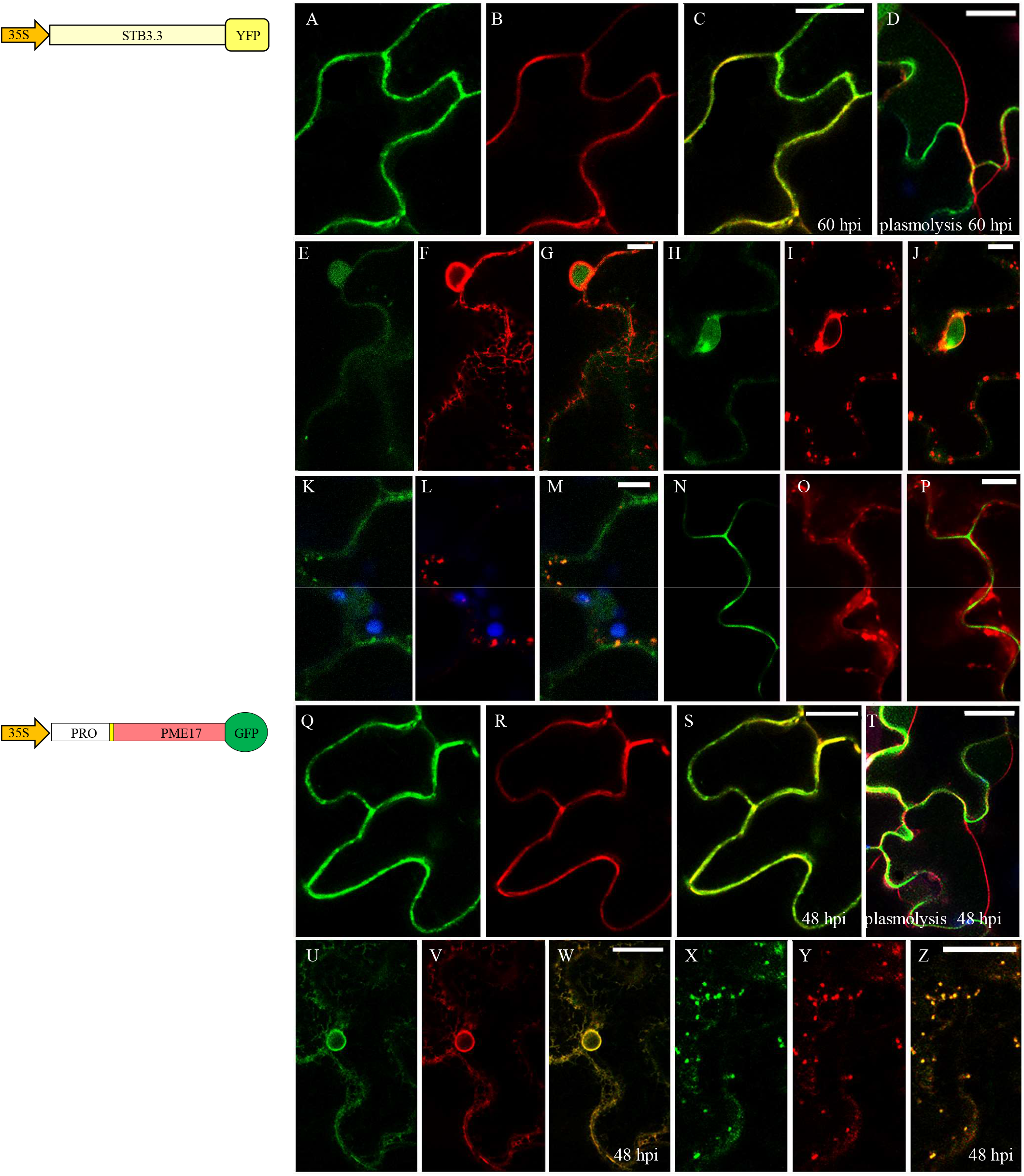
SBT3.3 and Pro-PME17 are delivered in the apoplast through different protein secretion pathways. Confocal microscopy images of epidermal cells of *Nicotiana tabacum* adult leaves showing the coexpression of SBT3.3-YFP (**A-P**) or Pro-PME17-GFP (**Q-Z**) with different markers of subcellular compartments (**B-C-, R-S:** XTH11-RFP, cell wall; **F-G, V-W** RFP-HDEL, endoplasmic reticulum; **I-J, Y-Z:** ST52-mCherry, Golgi stacks) at the indicated hours post infiltration (hpi). Plasmolyzed cells (1 M NaCl) expressing SBT3.3-YFP or Pro-PME17-GFP and the plasma membrane marker pm-rk show the green fluorescent cell wall and the retracted red fluorescent plasma membrane (**D, T**). Chlorophyll autofluorescence within the chloroplasts is depicted in blue. Scale bars = 20 μm (**A-C, Q-S**), 10 μm (**E-P**, **U-Z**), and 5 μm (**D, T**).

**Figure 9.**
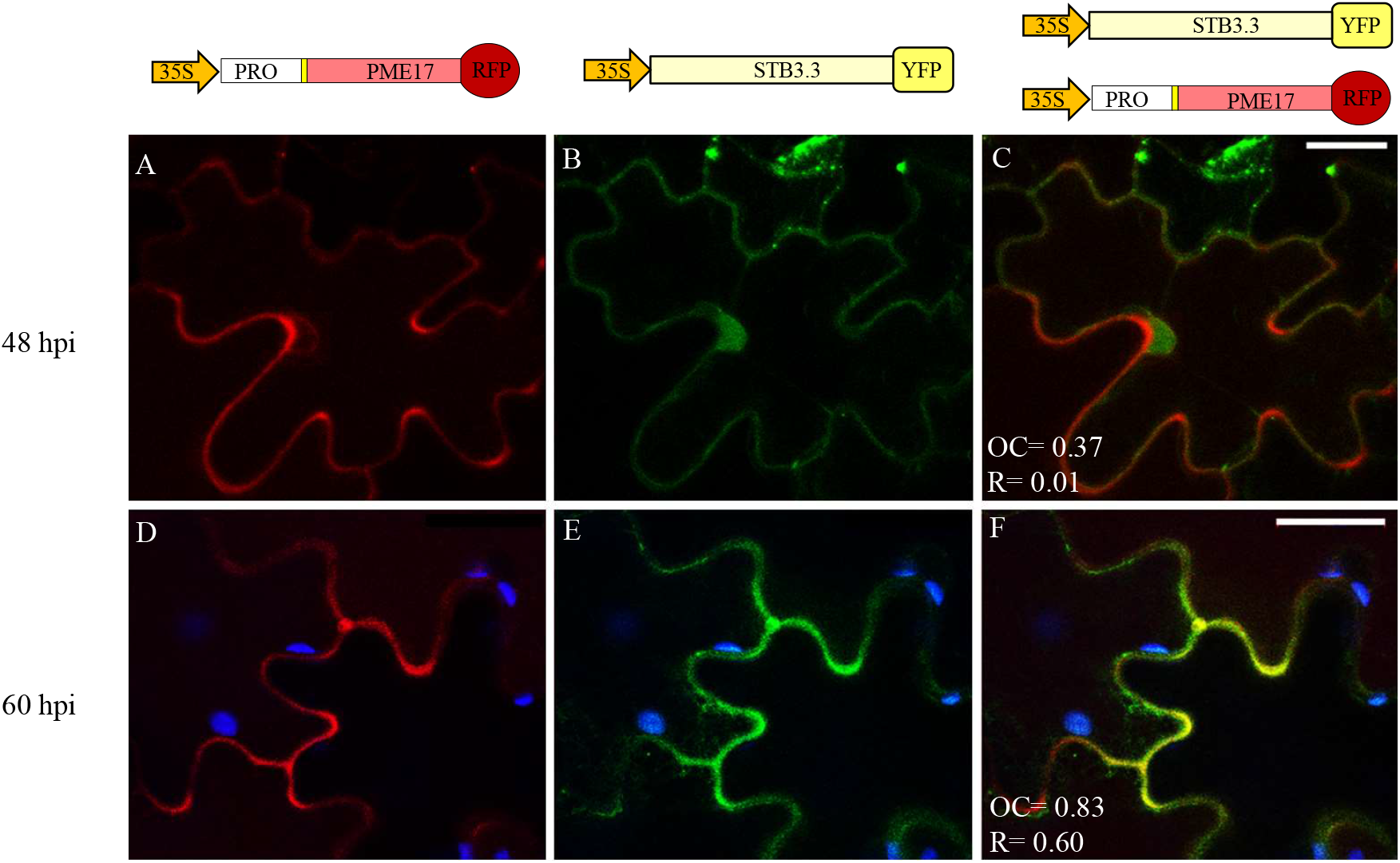
Pro-PME17 and SBT3.3 colocalize in the cell wall and their colocalization is favoured over time. Confocal microscopy images of epidermal cells of tobacco adult leaves co-expressing Pro-PME17-RFP (**A** and **D**) and SBT3.3-YFP (**B** and **E**) at the indicated hours post infiltration (hpi). Merged images showing the co-localization of SBT3.3-YFP and Pro-PME17-RFP in the cell wall (**C** and **F**). Overlap coefficient (OC, after Manders) and R (Pearson’s correlation coefficient) values are reported in each merged image. Chlorophyll autofluorescence within the chloroplasts is depicted in blue. Scale bars = 20 μm.

## DISCUSSION

Plants exploit a great and local increase of PME activity in response to pathogens with different lifestyles (Bethke et al., 2014; Lionetti et al., 2014; Lionetti, 2015; Lionetti et al., 2017). *A. thaliana* induces the expression of specific PME isoforms during the attack of different pathogens. Among them, *Pro-PME17* is induced after the infection of a wide range of plant pathogens and can be considered a general biomarker of pathogenesis (Lionetti et al., 2012). *Botrytis cinerea* is an important fungal plant necrotrophic pathogen, that kills plant tissue prior to feeding using a range of toxic molecules to destroy host cells (Williamson et al., 2007; Dean et al., 2012). Pro-PME17 was recently involved in triggering PME activity and in resistance against *B. cinerea* (Del Corpo et al., 2020). The Pro region acts as an intramolecular inhibitor of Pro-PME17 activity. Here, intriguingly we note that all the PME isoforms induced in *A. thaliana* during *B. cinerea* infection are organized as zymogens in which the catalytic region is preceded by an N-terminal Pro region. The structural similarity revealed between the amino acid sequences of the Pro regions of these PMEs and the independent immunity-related PMEIs (AtPMEI10 and AtPMEI11) (Lionetti et al., 2017; Del Corpo et al., 2020) support their possible role as intramolecular inhibitors of the specific PMEs in each specific Pro-PME cluster. We therefore attempted to identify which possible proteases could be involved in the activation of PME activity against microbes. Different SBTs were associated to the processing of PMEs in plant physiology (Schaller et al., 2018). Most SBTs are targeted to the CW, where they contribute to the control of CW properties during growth and development. The *Arabidopsis* SBT3.5 was previously found to process Pro-PME17 releasing PME activity during root growth in *Arabidopsis* seedlings (Senechal et al., 2014a). SBT1.7 modulates PME activity in the mucilage release from *Arabidopsis* seed coats (Rautengarten et al., 2008). AtSBT6.1 is likely to be involved in maturation of a pectin methylesterase (Wolf et al., 2009). The evidence that the precursors of bZIP transcription factors, RALF23 and GOLVEN1 are further substrates of AtSBT6.1 and that most SBTs are synthesized as inactive precursors could suggest that SBTs isoform could be part of an activation cascade, similar to that observed for caspases during plant programmed cell death events in mammals (Vartapetian et al., 2011; Reichardt et al., 2018; Paulus and Van der Hoorn, 2019).

Although SBTs were also associated to plant immunity their molecular substrates and their function remains largely unknowns (Figueiredo et al., 2014; Schaller et al., 2018; Godson and van der Hoorn, 2021). The tomato subtilisin-like protease SlSBT3 contributes to insect resistance in tomato. Although SlSBT3 was found to affect PME activity and the level of pectin methylesterification, these effects were unrelated to its role in herbivore defense (Meyer et al., 2016). Also, AtSBT5.2 and AtSBT3.14 were involved in different plant-pathogen interactions but their functions appear unrelated to PME activity (Lozano-Torres et al., 2014; Serrano et al., 2016). Here, we explored on the possible role of SBTs as post-transcriptional regulators of PME activity in *Arabidopsis* immunity against the necrotrophic fungus *B. cinerea*. Exploiting publicly available microarray data, we selected two genes, *SBT3.3* and *SBT3.5* co-expressed with *Pro-PME17* in *Arabidopsis* during *Botrytis* infection. SBT3.3 was previously discovered as a extracellular subtilase involved in the activation of *Arabidopsis* immunity against *Pseudomonas syringae* and *Hyaloperonospora arabidopsidis* (Ramirez et al., 2013) although its substrate remained unknown. Here, we demonstrated that *SBT3.3* and *SBT3.5* are induced in *Arabidopsis* leaves challenged with *Botrytis. SBT3.3* exhibited a higher fold-induction respect to SBT3.5. Interestingly, the expression of both *SBT* genes is also triggered in *Arabidopsis* against several pathogens. Moreover, *SBT3.3* and *SBT3.5* are induced by different bacterial and fungal elicitors of plant defense responses. SBT3.3 is specifically dedicated to plant defense because it is induced only during the infection process. However previous evidence indicates that SBT3.3 expression respond very rapidly to H2O2 and it enhances the activation of mitogen-activated protein kinases (MAPKs) following microbial attack (Ramirez et al., 2013). We detected SBT3.3 as a component of OGs signaling. All these evidence points to the involvement of SBT3.5 and, particularly of SBT3.3, as contributors in the activation of the earliest signaling events of PTI against *Botrytis* and several other pathogens.

The possible role of SBT3.3 and SBT3.5 as PME activators in *Arabidopsis* immunity against *B. cinerea* infection was explored. *sbt3.3-1*, *sbt3.3-2, sbt3.5-1*, *sbt3.5-2* mutants showed a significant higher susceptibility to the fungus when compared with WT. The significant lower induction of PME activity observed in *B. cinerea*-infected *sbt3.3* and *sbt3.5* mutants respect to WT plants indicates that SBT3.3 and SBT3.5 are involved in the release of mature active PMEs to contrast the microbial invasion. The susceptibility- and PME-related effects are more marked in the *sbt3.3* mutant respect to *sbt3.5* mutants suggesting that SBT3.3 can have a major contribution on the total PME activity expressed in *Arabidopsis* against the fungus. This also suggests that the two SBTs analyzed in this study can target different Pro-PME isoforms. The involvement of multiple SBT-Pro-PME pairs is also suggested by the observed variability in the processing motifs of all the PME isoforms involved in *Arabidopsis-Botrytis* interaction. We revealed a significant higher induction of PME activity and a higher resistance to *Botrytis* in infected *SBT3.3-OE1* and *SBT3.3-OE2* leaves respect to untransformed plants. Overall, our evidence indicates that SBT3.3 and SBT3.5, by processing different Pro-PME isoforms, contribute to trigger PME activity to contrast *B. cinerea* infection. The absence of growth abnormalities and alteration of monosaccharide composition in the *sbt* mutants and SBT3.3OE lines indicate that these SBTs are mostly involved in plant immunity processes in leaves of adult plants.

The low level of CW methylesters detected in SBT3.3OE lines indicate that SBTs, by timely activating PMEs early at the point of fungal penetration, can favor the increase of charged carboxyl groups forming a higher amount of Ca^2+^-mediated pectin crosslinks to reinforce the CW structure and the resistance to the microbe. Moreover, the SBT3.3-related pectin structural reinforcement is also accompanied by the stimulation of an immune response signaling. The higher induction of *WRKY33, PAD3, CYP81F2* and *WAK2* genes detected in *SBT3.3-OE1* plants respect to WT plants identifies a more robust immune response as mechanism contributing to the improved resistance observed in the transgenic plants. The induction of *WRKY33, PAD3* and *CYP81F2* by *SBT3.3* overexpression suggests the involvement of Camalexin and indole Glucosinolate accumulation. The evidence that WAK2 is highly expressed in transgenic plants suggest also the improvement of cell wall surveillance (Kohorn et al., 2014). Interestingly, WAK2 was demonstrated to require PME-mediated pectin de-methylesterification to activate OGs-dependent stress responses. We can hypothesize that SBT3.3 activates PME activity to favour WAK2 binding, and WAK2-OGs mediated stress signalling. Considering that OGs induce the expression of *SBT3.3*, *WRKY33*, *PAD3* and *CYP81F2* and that a *SBT3.3* self-activation was previously reported (Denoux et al., 2008; Ramirez et al., 2013), a positive feedback loop circuit based on PME activity, could be mounted to strengthen the activation of OGs-mediated defense responses. In addition, SBTs-dependent PME activity could also favour the binding and the functions of other PRR, like WAK1 and FERONIA, known to preferentially bind de-methylesterified crosslinked pectins (Decreux and Messiaen, 2005; Feng et al., 2018; Guo et al., 2018; Lin et al., 2018). Future efforts will be needed to explore on these hypotheses. All these observations indicate SBTs and PME activities exploit a cooperation between CW integrity maintenance and PTI signalling to regulate defense responses after a CW damage (Engelsdorf et al., 2018).

Although the specific pairs of SBT3.3, SBT3.5 and Pro-PMEs involved in plant immunity remain to be identified, we found that Pro-PME17 and SBT3.3 are both delivered in the apoplast although with different secretion pathways. While Pro-PME17 follows a conventional secretion pathway, SBT3.3 undergoes an unconventional route since bypass the Golgi, is BFA insensitive and colocalizes with Exo70E2, a subunit of the exocyst complex specifically identified in the EXPOs involved in unconventional exocytosis. To date, SAMS2, involved in lignin methylation and XTH29, an endotransglucosylase/hydrolases, are the two cell wall proteins reported to be secreted as EXPO cargo ((Wang et al., 2010; De Caroli et al., 2021). As previously proposed, EXPOs-mediate secretion emerges as a toolkit exploited by plants to ensure a dynamic remodelling of the cell wall structure able to adapt to changing environmental and growing conditions (de la Canal and Pinedo, 2018). Moreover, Pro-PME17 reaches the CW before SBT3.3 and their colocalization in the apoplast increase over time. These spatio-temporal separations could avoid a premature PME activation and pectin de-methylesterification during the secretion int the CW. Plants can accumulate Pro-PMEs clusters as soluble and inactive receptors in the apoplast. At the right time, SBTs could be delivered in an apoplast already enriched in the inactive Pro-PMEs, to unblock the inhibitory activity exerted by the Pro region in a compartment-specific manner, rapidly derepressing PME activity. This mechanism could lead to a faster induction of PME activity at the point of pathogen penetration to amplify the early immune signalling underlying the defense response. The speed of a response, particularly to biotic stress, can be crucial for the survival of the sessile plant. In a similar mechanism, known as ectodomain shedding, the proteolytic cleavage of extracellular domains of cell surface proteins resulting in the activation of a variety of normal and pathological signalling processes in animals including growth factor signalling, cell adhesion, inflammation and cell survival (Lichtenthaler et al., 2018). This proposed SBTs function would be also in line with the de-repression that speeds the responsiveness of hormonal transcription networks in plants (Rosenfeld et al., 2002). It is important to remember that also SBTs are Pro-peptides, in which the PRO region serves both as intramolecular chaperones required for enzyme folding and as inhibitor of the mature proteases (Hohl et al., 2017). All this evidence suggests the existence of mechanisms that accumulate multiple inactive molecular tools at the cell surface to be turned on at the right time to amplify a cell wall integrity signalling leading to cell solution to contrast pests. Many efforts are still required for the identification of the factors underlying these mechanisms although the pH variation in the apoplast also important during microbe invasion seems to be involved (Janzik et al., 2000; Geilfus, 2017).

Interestingly, SBT3.3 was involved in the physiological state called “priming”, a sensory status able to induce a resistance faster and stronger respect to the same stimulus received before (Ramirez et al., 2013). SBT3.3 expression mediates the activation of chromatin remodelling of defence genes including a self-activation, suggesting a positive feedback loop circuit to potentiate the defence response and keep cells in a sustained sensitised mode. Our observation on the localization of SBT3.3-YFP in the nucleus is consistent with this proposed role for this SBT. It was previously hypothesized that “priming” could involve the accumulation of inactive proteins that becomes operative to initiate a signal amplification leading to a faster and stronger activation of defense responses (Bruce et al., 2007; Conrath, 2011; Ramirez et al., 2013). The processing of Pro-PMEs by SBT3.3, timely accumulated, thanks to a spatial and temporal separation in the secretion of the two proteins in distinct subcellular compartments, could be part of the immune priming processes. Two PME-related DAMPs, OGs and MeOH are known to trigger defensive priming in plant immunity (Dorokhov et al., 2012; Komarova et al., 2014; Gamir et al., 2021; Giovannoni et al., 2021). The possibility that SBTs, trigger PMEs to assist and reinforce the OGs and MeOH signalling in priming is an interesting hypothesis to be investigated. It is conceivable that the proteolytic cleavage of other Pro-proteins by these and other SBTs could participate to trigger the immune response.

In conclusion, this study brings new insights into the molecular mechanisms underlying the activation of PME related immunity against necrotrophs increasing our understanding of the complexity of the interactions between plants and microbes. A switch on mechanism, mediated by SBTs for fine tuning PME activity and pectin integrity signaling, emerges providing knowledge about how plants modulate immunity. We identify Pro-PMEs as the target substrate processed by SBT3.3 indicating how this host factor mediates immune prime activation. Future studies will be useful to identify the specific SBT-Pro-PME pairs and to quantify the contribution of PME activity to priming. Activation of Pro-PMEs and plant immunity by SBTs processing becomes an effective genomic tool to engineer durable, broad-spectrum plant disease resistance and to avoid a growth-defense trade-offs.

## Materials And Methods

### Plant, Bacteria and Fungus growth conditions

*Arabidopsis thaliana* Wild-type Columbia (Col-0), *sbt* mutants and SBT3.3-OE seedlings were grown in solid MS/2 medium [Murashige and Skoog Medium including vitamins (2.2 g L^-1^), 1% sucrose, 0.8% plant agar, pH 5.7] in a controlled environmental chamber maintained at 22°C and 70% relative humidity, with a 16h/8h day/night cycle (PAR level of 100 μmol m^-2^s^-1^). Adult plants were transferred on soil in a growth chamber at 22°C and 70% relative humidity under a 12 h/12h day/night cycle (PAR level of 120 μmol m^-2^s^-1^). *Sbt* mutants used include SALK_086092 (At1g32960; *sbt3.3-1*), SALK_107460 (At1g32960; *sbt3.3-2*), SAIL_400_F09 (At1g32940; *sbt3.5-1*) and GABI_672C08 (At1g32940; *sbt3.5-2*). All mutants are in Col-0 background and are T-DNA insertion lines obtained from the Nottingham *Arabidopsis* Stock Centre (NASC). *Nicotiana tabacum cv* SR1 seeds were sterilized using commercial grade bleach solution in the presence of 0.1% (v/v) for 15 min at room temperature and then washed three times for 5 min with distilled water before sowing. The seeds were germinated in a growth room at 25°C and 30% relative humidity under a 16h/8h day/night cycle (photosynthetic photon flux of 50 μmol m^-2^s^-1^). Three- to four-week-old plants from MS medium were transferred on soil and grown in a controlled-environment at 25°C and 40% relative humidity under a 16h/8h day/night cycle (photosynthetic photon flux of 120 μmol m^-2^s^-1^). *Escherichia coli* TOP10 cells was grown in LB agar plate (Luria-Bertani: tryptone 1%, yeast exctract 0.5%, NaCl 1%) containing ampicillin 50μg/ml and incubated overnight at 37°C. *Agrobacterium tumefaciens* (Strain GV3110) was grown in LB medium (Luria-Bertani: tryptone 1%, yeast exctract 0.5%, NaCl 1%) containing appropriate antibiotics (Rifampycin 100μg/ml and Spectomycin 100μg/ml or Kanamycin 50μg/ml) and incubated for 2 days at 28°C. *Botrytis cinerea* strain SF1 (Lionetti et al., 2007) was grown for 20 days on Malt Extract Agar at 20 g L^-1^ with Mycological Peptone at 10 g L^-1^ (MEP) at 23°C and 70% relative humidity in the dark before conidia collection.

For seed germination analysis, *A. thaliana* WT, *sbt* mutants and SBT3.3-OE seeds were washed in 2 mL of isopropanol for 30 seconds followed by washing in 2 mL of sterile water for 3 min in slow agitation. Seeds were treated with 2 mL of sterilization solution (400 μl NaClO, 1.6 ml sterile water) for 5 min in slow agitation, followed by 4 washing steps in 2 ml of sterile water. For seed germination assay, triplicate sets of 60–70 sterilized seeds for each genotype were sown in Petri dishes on solid

MS/2 medium in a controlled environmental chamber maintained at 22°C and 70% relative humidity, with a 16h/8h day/night cycle (PAR level of 100 μmol m-2s-1). Seeds germination rate, as the number of germinated seeds divided by the total amount of total seeds, were scored at 24, 48 and 72 hours.

### Genotyping of T-DNA insertion mutants

Genomic DNA was extracted using RBC Bioscience Genomic DNA Extraction Kits, according to the manufacturer’s instructions, from leaves of 4-week-old *Arabidopsis* wild-type and mutant plants and subjected to a PCR-based screening using sequences of primer, described in Supplemental Table S1, generated with Primer3 (https://primer3.ut.ee/). Taq Kapa DNA Polymerase (Kapa Biosystem) was used at 1 units/30 μl of assay. Amplification was carried out in presence of dNTPs (0.2 mM), 2.5 mM MgCl2 and specific primers (10 μM) in the 1x buffer provide by the supplier. Conditions for amplification were: 95°C for 3 minutes; 36 cycles of amplification: 95°C for 30 sec, 60°C for 30 sec, 72°C for 60 sec, and a final extension of 72°C for 1 min. PCR products were separated by agarose gel electrophoresis and visualized by ethidium bromide staining. Plants homozygous for the mutations were propagated and used for all subsequent experiments.

### Stable transformation of SBT3.3-OE *Arabidopsis* plants

Flowering *Arabidopsis* plants were transformed by floral dipping for 15 seconds in 200 mL of the culture medium with the inoculum of *Agrobacterium tumefaciens* C58 carrying the construction of interest. The plants were subsequently covered for 1 day to promote the infection. The culture medium for immersion was composed of MS 2.2 g/L and sucrose 5% (p/v), supplemented with Silwet 0.005% and BAP 0.05 ug/mL. Subsequently, the selection of homozygous transgenic plants for the *SBT3.3- YFP* construction was carried out by means of selection with BASTA herbicide.

### Elicitor Treatment and Gene expression analysis

*B. cinerea*-infected or Mock-inoculated leaves were frozen in liquid nitrogen and homogenized using the mixer mill MM301 (RETSCH) and using inox beads (5 mm diameter) for about 1 min at 30 Hz, and total RNA was extracted with Isol-RNA Lysis Reagent (5’-Prime) according to the manufacturer’s instructions. RNA was treated with RQ1 DNase (Promega), and first-strand complementary DNA (cDNA) was synthesized using ImProm-II reverse transcriptase (Promega). Quantitative Reverse Transcription PCR analysis was performed using a CFX96 Real-Time System (Bio-Rad). One microliter of cDNA (corresponding to 50 ng of total RNA) was amplified in 20 μL of reaction mix containing 1× Go Taq qPCR Master Mix (Promega) and 0.5 μM of each primer described in Supplemental Table S2. Conditions for amplification were: 95°C for 2 minutes; 46 cycles of amplification: 95°C for 15 sec, 58°C for 15 sec, and a final extension of 72°C for 15 sec. Expression levels of each gene, relative to the UBIQUITIN5 (UBQ5) gene, were determined using a modification of the Pfaffl method (Pfaffl, 2001). Primer sequences were generated with Primer3 software (https://primer3.ut.ee/). Seedlings of 10-days-old *Arabidopsis* cultured in liquid MS/2 medium were treated for 1h with OG_s_ (40 μg mL^-1^) or water as a control, collected and processed for RNA extraction as described above.

### *Arabidopsis thaliana* infection with *Botrytis cinerea*

Conidia of *B. cinerea* were harvested by washing the surface of the mycelium with sterile distilled water. Conidia suspensions were filtered to remove residual mycelium and the conidia concentration was determined using a Thoma chamber. Fully developed leaves from 4-week-old *A. thaliana* plants were infected with 1×10^6^ conidia mL^-1^ incubated in Potato Dextrose Broth (PDB) at 24 g L^-1^. Six droplets of spore suspension (5 μL each) were placed on the surface of each leaf. Mock-inoculation was performed using PDB. Plants were incubated at 24°C with a 12h/12h day/night photoperiod. The lesion size produced by *B. cinerea* was evaluated as an indicator of susceptibility to the fungus.

### Determination of PME activity, fungal diffusion and H_2_O_2_ accumulation

Total protein extracts were obtained by homogenizing uninfected and infected *Arabidopsis* leaves in the presence of 1M NaCl, 12.5 mM Citric Acid, 50 mM Na2HPO4, 0.02 % Sodium Azide, protease inhibitor 1:200 v/v (P8849, Sigma), pH 6.5. Each sample was resuspended in 500 μL of protein Extraction Buffer. The homogenates were shaken for 3 h at 4°C, centrifuged at 13.000 x g for 15 min, and the supernatant (400 μL) collected. Protein concentration was determined in the supernatants using Bradford protein assay method (Bradford reagent, Sigma–Aldrich), a standard curve was performed using BSA (bovin serum albumine) at 1 mg mL^-1^. PME activity was evaluated by using PECTOPLATE assay (Lionetti, 2015): PECTOPLATE was prepared with 0.1% (w/v) of apple pectin (molecular weight range 30,000–100,000 Da; 70–75% esterification; 76282, Sigma-Aldrich, St. Louis), 1% (w/v) SeaKem^®^ LE agarose (Lonza, Basel, Switzerland, Catalog no: 50004E), 12.5 mM citric acid and 50 mM Na2HPO4, pH 6.5. Equal amounts of protein samples (2 μg of total protein in 20 μL) were loaded in each well of PECTOPLATE. Plates were incubated at 30°C for 16 h, and stained with 0.05% (w/v) Ruthenium Red (RR) (R2751; Sigma-Aldrich, St. Louis) for 30 min. The plates were de-stained by several washes with water and the area of the fuchsia-stained haloes, resulting from de-methylesterification of pectin and the area of inner unstained haloes, resulting from the hydrolysis of pectin in the gel, were measured with ImageJ software (Abramoff, Magalhes, and am 2004). Known amounts of commercially available PME from orange (Citrus spp.) peel (P5400; Sigma-Aldrich) was used in the PECTOPLATE to generate a standard curve used to calculate PME activity in the protein extracts.

For the detection of *B. cinerea* hyphae, detached leaves were incubated for 30 min in lactophenol Trypan Blue solution (water: glycerol: lactic acid: phenol [1:1:1:1] + Trypan Blue solution [1 mg mL^-1^; Sigma-Aldrich]). Thereafter, leaves were washed for overnight with gentle agitation in absolute ethanol to remove chlorophyll. Leaves were assayed for H2O2 accumulation employing 3,3’-diaminobenzidine staining as previously described (Daudi and O’Brien, 2012). Leaves were mounted in 50% glycerol and examined by light microscopy using Nikon eclipse E200 microscope. Images were taken with a Nikon Digital Sight DS-Fi1c camera

### Determination of Monosaccharide Composition of CW and MEOH quantification

Extraction of alcohol-insoluble residue, CW monosaccharide composition and the determination of CW methylesters were performed as previously described (Lionetti et al., 2017).

### SBT3.3-YFP and PME17-GFP/RFP Gene Constructs

For the SBT3.3-YFP overexpressing construct, a full-length cDNA for SBT3.3 was amplified by PCR using Pfu DNA polymerase (Stratagene, San Diego, CA) and specific primers, described in Supplemental Table S3, including Gateway adapters: BP SBT3.3 FW and BP SBT3.3 RV and recombined into pDONR207 using BP ClonaseMixII kit (Invitrogen). After sequencing, the construct was recombined with pB7FWG2 destination vector (YFP in C-terminal) using LR ClonaseMixII kit (Invitrogen) and introduced into *Arabidopsis* (*sbt3.3*) via Agrobacterium transformation.

For PME17-GFP/RFP overexpressing constructs, the full-length cDNA for PME17 was amplified by PCR using *TakaRa* Ex Taq DNA polymerase (TAKARA, Kyoto), and specific primers flanked by Gateway adapters, attB sequences, described in Supplemental Table S3. PCR amplified attB-flanked DNA was recombined into pDONR221 using BP ClonaseMixII kit (Invitrogen). Specific pDESTs, pK7FWG2 and pK7RWG2,0, were used for GFP and RFP fusion in C-terminus to obtain Pro-PME17- GFP and Pro-PME17-RFP, respectively. All constructs were checked by sequencing (Eurofins Genomics, https://eurofinsgenomics.eu/). The Gateway constructs (Pro-PME17-GFP, Pro-PME17-RFP, SBT3.3-YFP, SBT3.3-mCherry) and RFP-HDEL, ST52-mCherry, XTH11-RFP, pm-rk and Exo70E2-mCherry were introduced into *Agrobacterium tumefaciens* (Strain GV3110) and the agroinfiltration was performed in *Nicotiana tabacum* leaves as previously described (De Caroli et al., 2011).

### Transient transformation of *Nicotiana tabacum* leaves and confocal Laser Scanning Microscopy

Almost fully expanded *Nicotiana tabacum* 6-week-old leaves were infiltrated with a suspension of *Agrobacterium tumefaciens* bearing the relevant construct in 2mM Na3PO4; 50mM 2-(N-morpholine)- ethanesulphonic acid (MES); glucose 0.5%; acetosyringone 100μM; pH= 5.6 at an OD_600_=0.5. After 48 and 52 hours post infiltration (hpi) fluorescence was analyzed in infiltrated leaves by confocal microscopy. For co-infiltration, *Agrobacterium* cultures grown separately were adjusted to an OD_600_ = 0.5 and mixed prior to infiltration. For BFA (Sigma-Aldrich, http://www.sigmaaldrich.com) treatment, disks (1 cm diameter) of transformed leaves were treated by immersion in a solution of BFA at the final concentration of 100 μM, as reported by De Caroli et al. (2021). Plasmolysis was induced by incubating leaf disks in 1 M NaCl hypertonic solution for 10 min. Transiently transformed tobacco leaves were observed using a confocal laser scanning microscope LSM710 Zeiss (http://www.zeiss.com) mounting material in water as previously described (De Caroli et al., 2011). GFP and YFP were detected within the short 505–530 nm wavelength range, assigning the green color; RFP and mCherry were detected within 560–615 nm, assigning the red color. Excitation wavelengths of 488 and 543 nm were used. The laser power was set to a minimum and appropriate controls were made to ensure there was no bleed-through from one channel to the other. Images were processed using PHOTOSHOP7.0 (Adobe, https://www.adobe.com). For quantitative colocalization analyses, the overlap coefficient (OC, after Manders) and the Pearson’s correlation coefficient (R) were evaluated as previously reported (De Caroli et al., 2011).

### Bioinformatics tools

The Expression Angler tool of the Bio-Analytic Resource for Plant Biology (BAR, http://bar.utoronto.ca/welcome.htm) was used for co-expression analysis. E-plant (https://bar.utoronto.ca/eplant/) was used for to study developmental map and biotic stresses. Chromas was used to manage both nucleotide sequences and amino acids (https://chromas.software.informer.com/2.5/). Primer3 (https://primer3.ut.ee/) was used to design primers respectively for Gene expression analysis and Genotyping analysis. ImageJ was used to calculate the area of the haloes in PECTOPLATE assay (http://imagej.nih.gov/ij/download.html). DNAMAN software (Lynnon, BioSoft, Quebec, Canada, https://www.lynnon.com/) was used for amino acid sequence analyses. Genevestigator Meta-Analyzer Tools (www.genevestigator.ethz.ch/at/) was used to analyze *SBT3.3* and *SBT3.5* gene expression during infection of *Arabidopsis* with different pathogens and DAMPs and PAMPs treatments.

### Accession numbers

The genes used in this study have the following accession numbers: SBT3.3, At1g32960; SBT3.5, At1g32940; UBQ5, At3g62250; PAD3, At3g26830; CYP81F2, At5g57220; PME17, At2g45220; PMEI10, At1g62760; WAK2, At1G21270; At2g38470, WRKY33. *B. cinerea* β-tubulin, MG949129.1.

## Supplemental data

**Supplemental Figure S1.** Gene structure and alignment of amino acid sequences of Pro-PMEs induced in Arabidopsis during *B. cinerea* infection.

**Supplemental Figure S2.** *SBT3.3* and *SBT3.5* gene expression in *Arabidopsis* challenged with *Botrytis cinerea*.

**Supplemental Figure S3.** *SBT3.3* and *SBT3.5* gene expression during infection of *Arabidopsis* with different pathogens and after different DAMPs and PAMPs treatments

**Supplemental Figure S4.** Seed morphology and germination are not altered in *sbt3.3, sbt3.5* mutants and *SBT3.3*-OE lines.

**Supplemental Figure S5**. Arabidopsis *sbt3.3-1, sbt3.5-1* mutants and *SBT3.3-OE1* line are not defective in H2O2 accumulation against *Botrytis*.

**Supplemental Table S1.** Primers used for genotyping

**Supplemental Table S2.** Primers used for qRT-PCR

**Supplemental Table S3.** Primers used to obtain Pro-PME17 and SBT3.3 fluorescent constructs with Gateway system.

## Acknowledgments

We thank Marco Greco for technical assistance.

## Funding

The work was supported by Sapienza University of Rome, Grants RM120172B78CFDF2, RM11916B7A142CF1 and RG12117A898EABE0 to VL and AR12117A8A4A1ADC to DC and LV.

## Author contributions

V.L. conceived the research; G.P, P.V and L.V. developed the experimental designs and supervised the experiments; D.C., D.D.C, M.O.M and M.D.C performed the experiments; D.C., M.D.C, G.P, P.V. and V.L. analysed data; All authors contributed to discussing the results of this manuscript. V.L. wrote the article with contributions of all the authors

## Notes

### Competing Interest Statement

The authors have declared no competing interest.

### Summary of Updates

This revised version of the manuscript has been improved in both presentation of results and content, adding further findings.

