## Supplementary Figures and Tables for "Subtilases turn on pectin methylesterase activity for a robust apoplastic immunity"

**A**

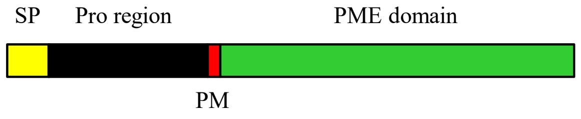


**B**

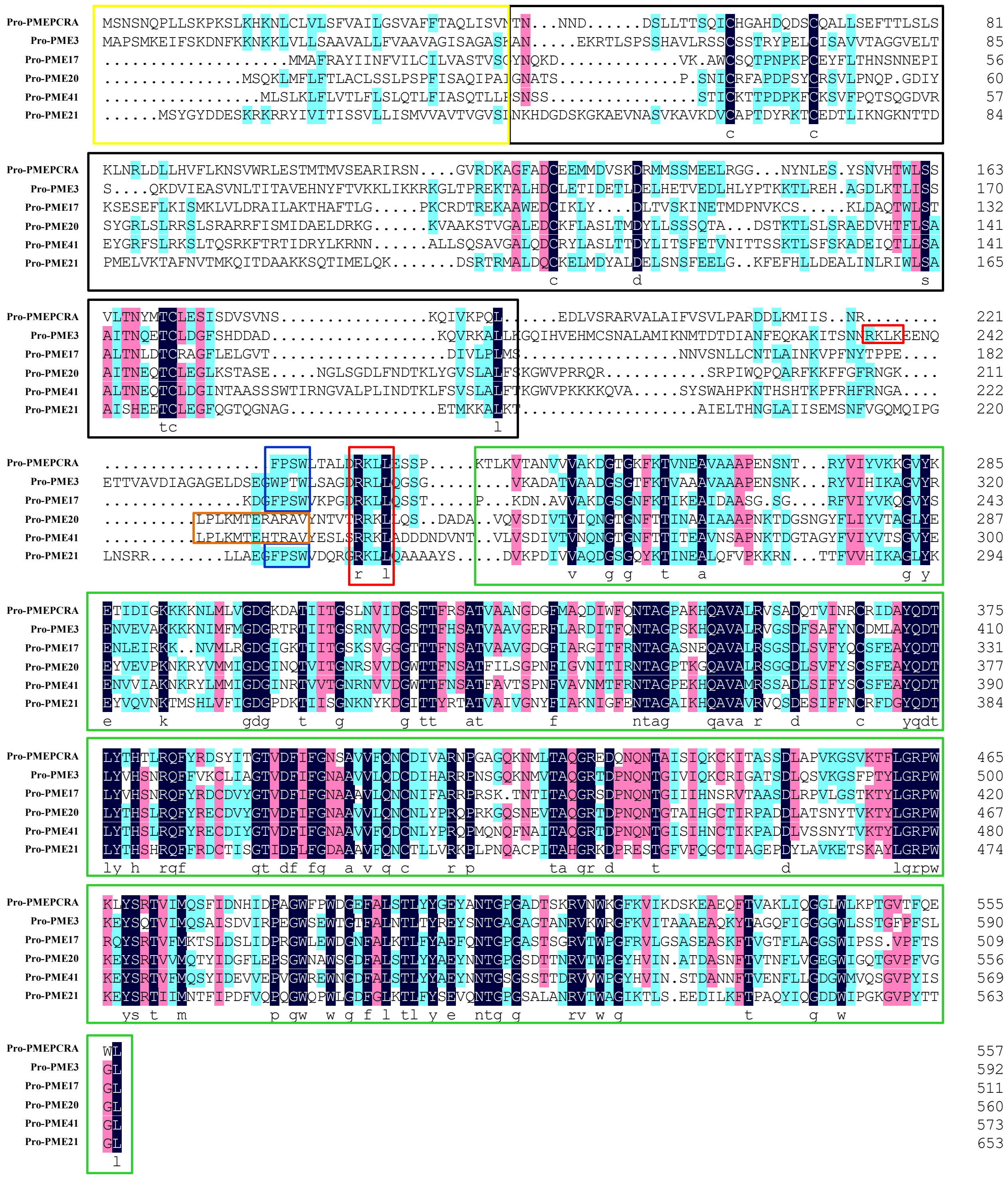


**Supplemental Figure S1. Gene structure and alignment of amino acid sequences of Pro-PMEs induced in Arabidopsis during *B. cinerea* infection. A)** Schematic representation of PME gene structure. **B)** Pro regions (black boxes) present the four conserved cysteine typically engaged in the formation of two disulfide bridges known to stabilize the structure in the independently-expressed PMEIs. A variability in the processing motif (PM; red box) was revealed. Other partially conserved motifs with an unknow function were also detected (blue and orange boxes). The PME catalytic domain (green box) is highly conserved. Highlight homology level: 100% black; >75% pink; >50% light blue. SP=Signal Peptide; PM=Processing Motif. DNAMAN software (Lynnon, BioSoft, Quebec, Canada) was used for amino acid sequence analyses.

**A**

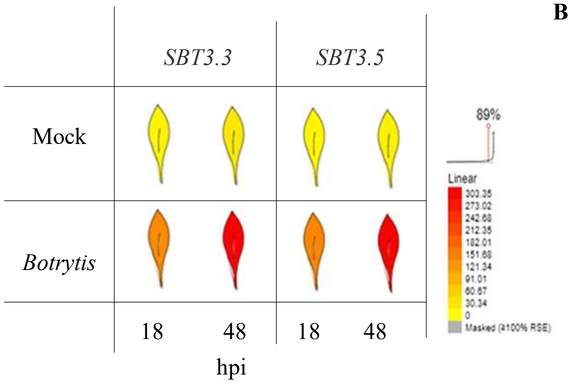


**B**

| Gene name | R-value |
|---|---|
| *Pro-PME17* | 1.000 |
| *SBT3.5* | 0.899 |
| *SBT3.3* | 0.594 |

**Supplemental Figure S2. *SBT3.3* and *SBT3.5* gene expression in *Arabidopsis* challenged with *Botrytis cinerea* A)** Expression and **B)** co-expression analysis were performed using Expression Angler tool of the Bio-Analytic Resource for Plant Biology (BAR, http://bar.utoronto.ca/welcome.htm). R-value= Pearson correlation coefficient.

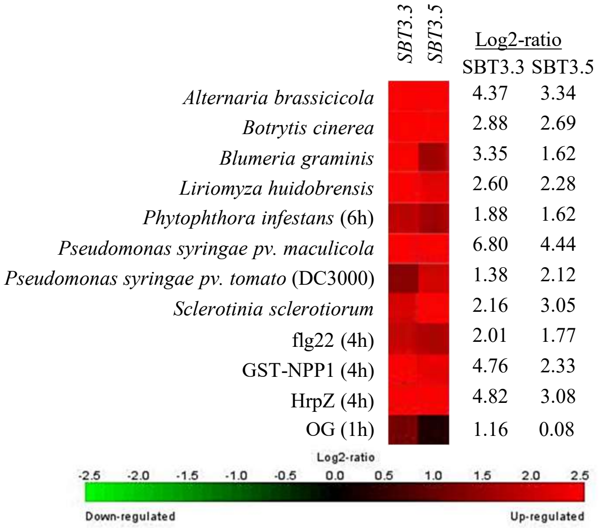


**Supplemental Figure S3. *SBT3.3* and *SBT3.5* gene expression during infection of *Arabidopsis* with different pathogens and after different DAMPs and PAMPs treatments** Expression of *SBT3.3* and *SBT3.5* in *Arabidopsis* leaves treated with different pathogens, the bacterial PAMP flg22 (flagellin), fungal PAMP GST-NPP1, HrpZ Effector and OGs. Data were analyzed using the Genevestigator Meta-Analyzer Tools (www.genevestigator.ethz.ch/at/). The specific expression level is indicated as *Log(2)*-ratio in treated plants relative to corresponding control. Filters used = Fold change ≥ 2; p-value ≤ 0.01.

**A**

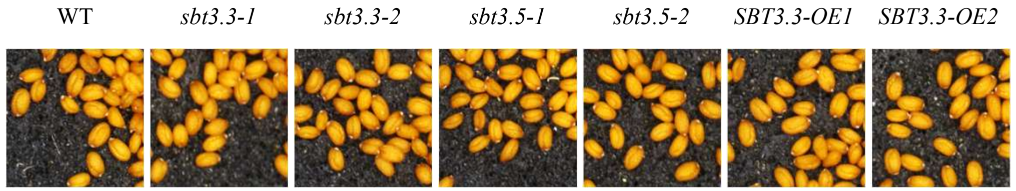


**B**

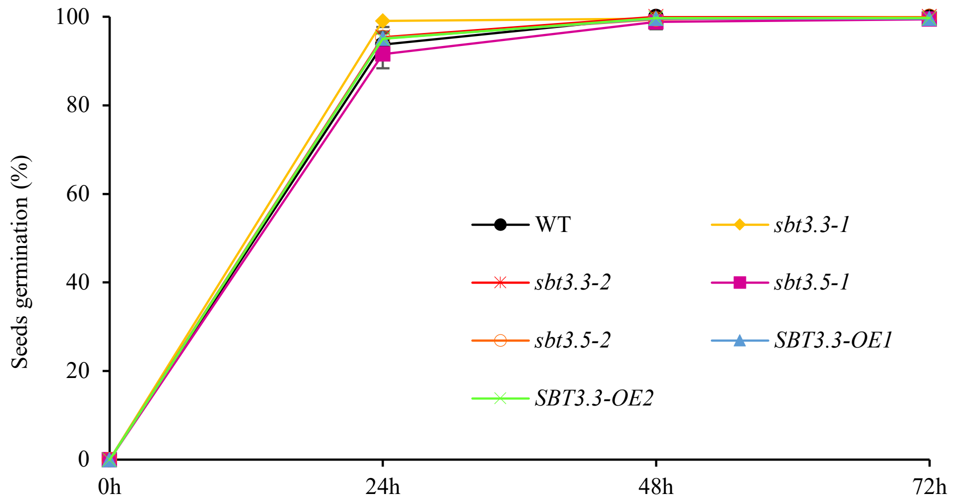


**Supplemental Figure S4. Seed morphology and germination are not altered in *sbt3.3*, *sbt3.5* mutants and *SBT3.3-OE* lines. A)** Representative images of s*bt* mutants, *SBT3.3-OE* and WT seeds. **B)** The germination percentage of mutants and WT seeds is calculated as the number of germinated seeds divided by the total amount of total seeds sown in the medium. Data represent mean ± SD of three replicates.

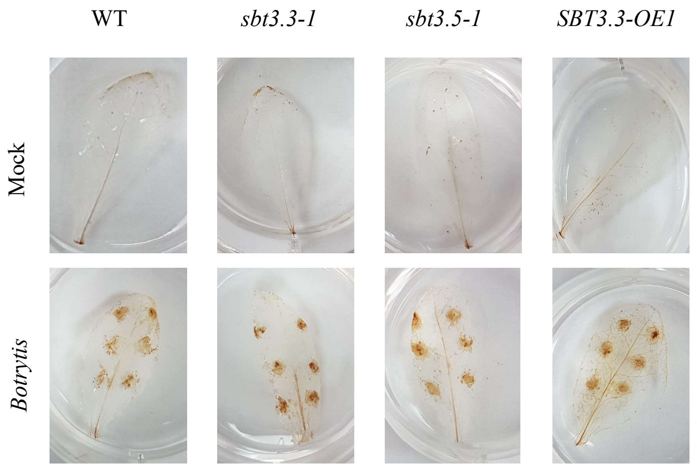


**Supplemental Figure S5. Arabidopsis *sbt3.3-1*, *sbt3.5-1* mutants and *SBT3.3-OE1* line are not defective in $H_2O_2$ accumulation against *Botrytis*.** Leaves at 24-hour post inoculation with Mock and *Botrytis* were stained with 3,3′-diaminobenzidine.

**Supplemental Table S1. Primers used for genotyping**

| AGI Code | Mutant name | T-DNA insertion line | Forward primer 5'-3' | Reverse primer 5'-3' |
|---|---|---|---|---|
| At1g32960 | *sbt3.3-1* | SALK_086092 | TCACACACACCTTGTCT TTGC | AAAACGCATGCGAACAT TTAC |
| At1g32960 | *sbt3.3-2* | SALK_107460 | CTTGGGGAGGAAGAAA AGATG | GAGGACCTAACTCGATG TCCC |
| At1g32940 | *sbt3.5-1* | SAIL_400_F09 | TTTAAATGGGCCTTAAA TCCG | CCAAGTTCGAGTTGTAGC GAG |
| At1g32940 | *sbt3.5-2* | GABI_672C08 | TTTAAATGGGCCTTAAA TCCG | GCTTGTGACTCGGTAAGC TTG |

Primers T-DNA 5'-3'

| | | |
|---|---|---|
| SALK | LBb1.3 | ATTTTGCCGATTTCGGAAC |
| SAIL | LB1 | GCCTTTTCAGAAATGGATAAATAGCCTTGCTTCC |
| GABI | o8474 | ATAATAACGCTGCGGACATCTACATTTT |

**Supplemental Table S2. Primers used for qRT-PCR**

| AGI Code | Name | Forward primer 5'-3' | Reverse primer 5'-3' |
|---|---|---|---|
| At3g62250 | *UBQ5* | CCGTGGTGGTGCTAAGAAGA | AGCTCCACAGGTTGCGTTAG |
| At1g32960 | *SBT3.3* | TTTTGCAGAAGGGTCATCTCG | GTCGTTGTAACCAGCAGAACA |
| At1g32940 | *SBT3.5* | CTCTACTTATGCTCCGCCGG | TAGTGACGGTTCGGGTGAGA |
| At3g26830 | *PAD3* | TCGCTGGCATAACACTATGG | TTGGGAGCAAGAGTGGAGT |
| At5g57220 | *CYP81F2* | TTTTCGGTTGTGTCCAGCTG | CAGGCATAAACTTCTCGGGC |
| At2g45220 | *PME17* | AAAGCAGCGTGGGAAGATTG | GCTCTAAGAACCCAGCTCGA |
| At1g62760 | *PMEI10* | CATATCTCACGGAGGTGGTC | CATTCGTGAGAGCAGCACTA |
| At1G21270 | *WAK2* | AACTGCCCATCTGGTTACCG | CTCTGTGTTCTTCCGGTGCT |
| At2g38470 | *WRKY33* | GAAACAAATGGTGGGAATGG | TGTCGTGTGATGCTCTCTCC |
| MG949129.1 | *B. cinerea β-tubulin* | TACCACCTGTCTCCGTTTCC | GCGGCCATCATGTTCTTAGG |

**Supplemental Table S3. Primers used to obtain Pro-PME17 and SBT3.3 fluorescent constructs with Gateway system.**

| Name | Forward primer 5’-3’ | Reverse primer 5’-3’ |
|---|---|---|
| **attB-** SBT3.3 | **GGGGACAAGTTTGTACAAAAAAGCAGGCTAT**ATGAGGAGTTTCAGATCATCCATTCTC | **GGGGACCACTTTGTACAAGAAAGCTGGGTA**TAATCCGTTCTCATCATAGTAGTTTTGCAG |
| **attB-** PME17 | **GGGGACAAGTTTGTACAAAAAAGCAGGCT**TCACCATGATGGCTTTTCGAGCTTA | **GGGGACCACTTTGTACAAGAAAGCTGGGTA**CTTGAGACCAGAAGTGAACGGCA |

AttB sequences are written in bold.
